# Dynamics and Regulatory Roles of RNA m^6^A Methylation in Unbalanced Genomes

**DOI:** 10.1101/2024.07.05.602246

**Authors:** Shuai Zhang, Ruixue Wang, Kun Luo, Shipeng Gu, Xinyu Liu, Junhan Wang, Ludan Zhang, Lin Sun

**Affiliations:** Key Laboratory of Cell Proliferation and Regulation Biology of Ministry of Education, College of Life Sciences, Beijing Normal University, Beijing 100875, China; Beijing Key Laboratory of Gene Resource and Molecular Development, College of Life Sciences, Beijing Normal University, Beijing 100875, China

**Author notes:** For correspondence (Lin Sun). These authors contributed equally to this work (Shuai Zhang, Ruixue Wang).

## Abstract

N^6^-methyladenosine (m^6^A) in eukaryotic RNA is an epigenetic modification that is critical for RNA metabolism, gene expression regulation, and the development of organisms. Aberrant expression of m^6^A components appears in a variety of human diseases. RNA m^6^A modification in Drosophila has proven to be involved in sex determination regulated by Sxl and may affect X chromosome expression through the MSL complex. The dosage-related effects under the condition of genomic imbalance (i.e., aneuploidy) are related to various epigenetic regulatory mechanisms. Here, we investigated the roles of RNA m^6^A modification in unbalanced genomes using aneuploid Drosophila. The results showed that the expression of m^6^A components changed significantly under genomic imbalance, and affected the abundance and genome-wide distribution of m^6^A, which may be related to the developmental abnormalities of aneuploids. The relationships between methylation status and classical dosage effect, dosage compensation, and inverse dosage effect were also studied. In addition, we demonstrated that RNA m^6^A methylation may affect dosage-dependent gene regulation through dosage-sensitive modifiers, alternative splicing, the MSL complex, and other processes. More interestingly, there seems to be a closely relationship between MSL complex and RNA m^6^A modification. It is found that ectopically overexpressed MSL complex, especially the levels of H4K16Ac through MOF could influence the expression levels of m^6^A modification and genomic imbalance may be involved in this interaction. We found that m^6^A could affect the levels of H4K16Ac through MOF, a component of the MSL complex, and that genomic imbalance may be involved in this interaction. Altogether, our work reveals the dynamic and regulatory role of RNA m^6^A modification in unbalanced genomes, and may shed new light on the mechanisms of aneuploidy-related developmental abnormalities and diseases.

## Introduction

Epigenetic modifications regulate gene expression in response to environmental changes and play important roles in the development of organisms and a variety of human diseases (Jaenisch and Bird, 2003; Lence et al., 2017). In addition to DNA and chromatin modifications, which are well studied, more than one hundred RNA chemical modifications have been identified in cells to date (Lee et al., 2014; Roundtree et al., 2017a). As a marker of post-transcriptional regulation, RNA modifications participate in almost all aspects of RNA metabolism (Roundtree et al., 2017a). N^6^-methyladenosine (m^6^A) is the most prevalent internal modification in many eukaryotic messenger RNAs (mRNAs) and long noncoding RNAs (lncRNAs) (Lee et al., 2014; Roundtree et al., 2017a; Yang et al., 2018), which widely affects RNA alternative splicing (Dominissini et al., 2012), export (Roundtree et al., 2017b), stability (Wang X et al., 2014), and translation (Meyer et al., 2015). RNA m^6^A modifications have been found to be enriched on the transcripts of genes that regulate development and cell fate specification (Dominissini et al., 2012; Meyer et al., 2012; Geula et al., 2015), and some m^6^A sites are regulated in a tissue- or disease-specific manner (Zaccara et al., 2019). In addition, the abnormal expression of m^6^A components is related to the tumorigenesis, proliferation, and metastasis of many types of cancers (Pinello et al., 2018; Ma et al., 2019). Therefore, it is of great significance to study RNA m^6^A methylation for revealing the mechanisms of gene expression regulation and human diseases.

At present, there are few studies on RNA m^6^A modification in Drosophila, possibly due to its relatively low abundance (m^6^A/A < 0.2%) (Haussmann et al., 2016), and the mutation of some m^6^A component genes will affect their viability and fertility (Hongay and Orr-Weaver, 2011; Haussmann et al., 2016; Kan et al., 2017). However, as a model organism with specific genetic and developmental advantages, Drosophila remains an excellent tool for studying the roles of epigenetic modifications in gene regulation, individual development, and disease process. Several components of the Drosophila m^6^A methyltransferase complex (Ime4, dMettl14, and fl(2)d constitute the core complex, with vir and nito acting as cofactors) and a m^6^A reader protein (Ythdc1) have been identified, all of which have homologues in mammals (Lence et al., 2017). Deletion of the major methyltransferase gene Ime4 or the reader Ythdc1 causes locomotion defects, and the splicing of sex-determining factor Sxl is affected (Haussmann et al., 2016; Lence et al., 2016; Kan et al., 2017). However, homozygous mutations in fl(2)d, vir, and nito were lethal, suggesting that these subunits have important functions other than methylation (Penn et al., 2008; Yan et al., 2015; Kan et al., 2017; Lence et al., 2017). RNA m^6^A modification has an obvious sexual dimorphism in Drosophila, and reduced m^6^A levels severely decreased the survival of females. It is thought to be due to the derepression of msl-2 caused by aberrantly spliced Sxl, which forms the Male Specific Lethal (MSL) complex that associates with the X chromosome (Haussmann et al., 2016).

Dosage compensation is a widespread phenomenon in unbalanced genomes (Lucchesi, 2018; Birchler and Veitia, 2021). The deletion or duplication of some chromosomes rather than the whole chromosome set leads to genomic imbalance, that is, aneuploidy (Orr et al., 2015). Aneuploid variation is usually detrimental to organisms (Birchler and Veitia, 2012; Orr et al., 2015; Birchler and Veitia, 2021), and is associated with developmental abnormalities, mental retardation, and various congenital defects (Williams et al., 2008; Huang et al., 2021; Sanchez-Pavon et al., 2021), possibly due to disorders in their gene expression systems (Prestel et al., 2010; Letourneau et al., 2014). Studies across species have pointed out that there is genome-wide trans modulation in aneuploidy, and the genes on the varied chromosomes are compensated to a certain extent, while genes located on the rest of the genome are mainly regulated in the opposite direction to the changes of chromosome numbers, which is known as the inverse dosage effect (Birchler and Veitia, 2012, 2021; Sun et al., 2013c; Hou et al., 2018; Shi et al., 2021). Histone modification (Zhang et al., 2021a), chromatin remodeling (Birchler, 2016), lncRNAs (Zhang et al., 2023) and microRNAs (Shi et al., 2022) have all been shown to play a role in genomic imbalance.

Because of the haploinsufficiency for X-linked genes, heterogametic individuals in organisms with XY sex determination systems could be regarded as analogous to aneuploidy (Disteche, 2016), including humans and Drosophila. Some studies have linked histone H4 lysine 16 acetylation (H4K16Ac) and non-coding roX RNAs to the dosage compensation of Drosophila (Park et al., 2010; Conrad et al., 2012). On the other hand, the compensation of X chromosome in human is thought to be regulated by lncRNA X-inactive specific transcript (XIST) (Jordan et al., 2019). Interestingly, in Drosophila, RNA m^6^A modification indirectly affects gene expression through Sxl and msl-2; while in human, m^6^A methylation of the key lncRNA XIST is necessary for the silencing of gene transcription on one of the female X chromosomes (Patil et al., 2016).

Moreover, most tumor cells have genomic instability and high levels of aneuploidy (Ben-David and Amon, 2020; Chiarle, 2021), and at the same time, m^6^A component genes are often aberrantly expressed in various cancers (Pinello et al., 2018; Ma et al., 2019).

An increasing number of studies have found that dosage-related effects in aneuploidy may be the integration of multiple modulations rather than through a single mechanism (Prestel et al., 2010; Birchler, 2016). The effects of genomic imbalance are complicated, and the model of dosage compensation and global gene regulation in Drosophila has been extended to autosomal aneuploidies and sex chromosome aneuploid metafemales where MSL complexes are not assembled (Sun et al., 2013b,c; Zhang et al., 2021b). In addition, genomic imbalance and trans regulatory mechanisms in other species such as maize, Arabidopsis, and humans have also been investigated (Hou et al., 2018; Raznahan et al., 2018; Shi et al., 2021; Yang et al., 2021; San Roman et al., 2023). To reveal the role of RNA m^6^A modification in unbalanced genomes, we studied the dynamic changes and regulatory functions of m^6^A methylation under genetic imbalance conditions using autosomal and sex chromosome aneuploid Drosophila maintained in our laboratory. Meanwhile, dosage-sensitive modifiers, differential alternative splicing events, the MSL complex and other factors that may mediate the relationships between RNA m^6^A modification and the dosage-related effects of aneuploidy were also investigated. In summary, we provided a comprehensive picture of RNA m^6^A methylation in unbalanced genomes.

## Results

### The responses of m A components under genomic imbalance

RNA m^6^A methylation is a reversible epigenetic modification, and its dynamic process is mediated by m^6^A methyltransferases (writers), demethylases (erasers), and m^6^A recognition proteins (readers) (Lee et al., 2014; Yang et al., 2018; Zaccara et al., 2019) (Figure 1A). We first detected the expression of m^6^A components in Drosophila larvae with karyotypes of normal diploids and trisomies that have an additional chromosome arm using RT-qPCR to determine whether RNA m^6^A dynamics are affected by correlative enzymes in unbalanced genomes (Figure 1B; Figure 1—figure supplement 1A,B). It was found that the transcription levels of most m^6^A writers and the major m^6^A reader are down-regulated in aneuploids compared with their respective sex-corresponding controls (Figure 1B). Unlike the other components, the cofactor vir of methyltransferase complex is up-regulated in trisomy 2L females (Haussmann et al., 2016; Lence et al., 2016; Kan et al., 2017). Because RNA m^6^A modification is enriched in the nervous system of Drosophila, the brains of aneuploid third instar larvae were also used to detect the expression of m^6^A components. As expected, a decreased expression trend of m^6^A components similar to that of the whole larvae was observed (Figure 1—figure supplement 1C).

**Figure 1.**
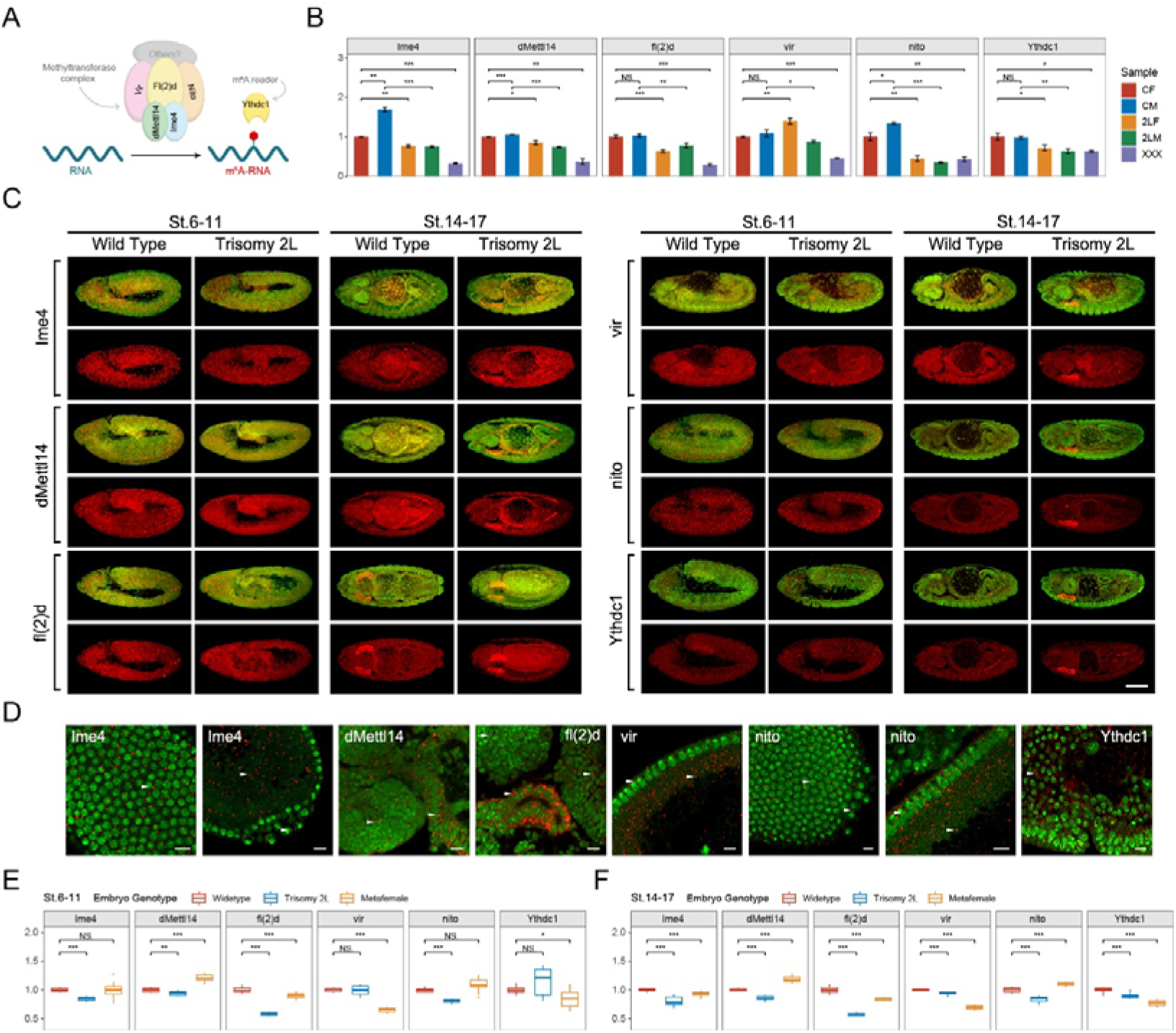
The responses of m^6^A methyltransferases and reader protein under the condition of genomic imbalance. (A) Schematic diagram of m^6^A components in Drosophila. (B) RT-qPCR analysis of mRNA levels of m^6^A methyltransferases and reader protein in third instar larvae of wildtype and trisomy Drosophila. CF, wildtype female control; CM, wildtype male control; 2LF, trisomy 2L female; 2LM, trisomy 2L male; XXX, metafemale; 2L, chromosome 2 left arm. Student’s t test *p < 0.05, **p < 0.01, ***p < 0.001. (C) Subembryonic distribution patterns of the transcripts of m^6^A components in wildtype and trisomy 2L Drosophila. The names of the genes were shown in the left of the pictures; the genotypes and stages were shown above. Red, probes; green, DAPI. Scale bar, 100 μm. (D) Subcellular localization of probe signals. Probe name was written in the corner of each picture. Red, probe; green, DAPI. Arrowheads indicate the foci of probe signals. The tissue types are (1) blastoderm nuclei; (2) yolk plasm and pole cells; (3) brain and midgut; (4) salivary gland and midgut; (5) blastoderm nuclei and yolk cortex; (6) blastoderm nuclei and pole cells; (7) blastoderm nuclei and yolk cortex; (8) germ band. Scale bars, 10 μm. (E,F) The expression levels of m^6^A component genes in stage 6-11 (E) and stage 14-17 (F) represented by relative fluorescence intensity of probes compared with DAPI signals. The expression of wildtype embryos was set as one. Sample size = 10. Student’s t test *p < 0.05, **p < 0.01, ***p < 0.001.

Previous studies have pointed out that the m^6^A methylomes have temporal and spatial specificity in the development process of organisms, especially for embryonic development and cell differentiation (Meyer and Jaffrey, 2014; Wang Y et al., 2014). Aberrant expression of m^6^A components may cause defects in embryogenesis and even early embryonic lethality (Zhong et al., 2008; Wang Y et al., 2014; Geula et al., 2015; Zhao et al., 2017). Therefore, we designed probes of m^6^A components (Figure 1—figure supplement 1D,E) to examine the mRNA expression and localization patterns during early embryonic development of aneuploid Drosophila using high-resolution TSA-FISH (Lecuyer et al., 2007; Jandura et al., 2017). The results showed that the subembryonic distribution patterns of the five components of m^6^A methyltransferase complex and one m^6^A reading protein are similar in wildtype and aneuploidies (Figure 1C,D; Figure 1—figure supplement 2A-F). At the blastoderm stage (stage 1-5), the probe signals of m^6^A components are widely distributed in the surface cell layer, yolk plasma, yolk cortex, and show a pattern of basal enrichment (Figure 1—figure supplement 2A-F). For the gastrulae at stage 6-11, the transcripts of m^6^A components are mainly located in head, amnioproctodeal invagination, germ band, and as in previous studies (Lence et al., 2016), and show an enrichment in neuroectoderm (Figure 1C; Figure 1—figure supplement 2A-F). At later stages of embryonic development (stages 12-13 and 14-17), the probes are distributed in brain, ventral nerve cord, midgut, and salivary gland (Figure 1C; Figure 1—figure supplement 2A-F).

By observing the subcellular localization of the probe signals of m^6^A components at higher magnification, it can be found that most of the mRNAs of these genes have nuclei-associated localization patterns (Figure 1D; Figure 1—figure supplement 2A’-F’). In the early and late stages of embryonic development, the probe signals of Ime4, dMettl14, fl(2)d, vir, and Ythdc1 form dense small foci near the nucleus, which is a perinuclear distribution (Figure 1D; Figure 1—figure supplement 2A’-F’). For nito, in addition to the perinuclear signals, there is also an obvious signal of intranuclear localization during early embryogenesis, which is manifested as one or two small foci in the blastoderm nucleus (Figure 1D; Figure 1—figure supplement 2E’).

Although there seems to be no difference in the localization of the transcripts of m^6^A components that we detected in aneuploidy and wildtype embryos, the expression levels of these genes, as determined by relative fluorescence intensity, are significantly changed (Figure 1E,F; Figure 1—figure supplement 2A”-F”). Except for the irregular fluctuations at early embryonic stages, the expression levels of most m^6^A components in aneuploids are lower than those in wildtype at more mature stages (Figure 1E,F; Figure 1—figure supplement 2A”-F”), which is similar to the trend we detected in third instar larvae (Figure 1B; Figure 1—figure supplement 1C). The transcripts of m^6^A methyltransferase and reader protein have specific subembryonic and subcellular localization patterns in the development of Drosophila embryos, which may be a mechanism to regulate cellular functions, and ensure appropriate cell growth and differentiation (Lecuyer et al., 2007).

### Genome-wide mapping of RNA m A methylation in aneuploid Drosophila

Subsequently, we detected the global levels of RNA m^6^A methylation in wildtype and aneuploid Drosophila larvae to determine whether m^6^A abundance is altered under the condition of genomic imbalance due to modulation by transmethylases. The overall abundance of m6A was represented by the m^6^A/A ratio in total RNA (Figure 2A). It was found that the m^6^A abundance of wildtype males is higher than that of females, but both are lower than 0.2%, which is consistent with the finding that m^6^A levels in Drosophila are relatively low (Haussmann et al., 2016; Lence et al., 2016). Females with triple chromosome 2 left arms (2L) and metafemales with triple X chromosomes have significantly higher m^6^A abundances than diploid females, whereas there is no significant difference between trisomy 2L males and wildtype males (Figure 2A). Therefore, genomic imbalance can affect the m^6^A methylation status to some extent, and this epigenetic modification is different between males and females. However, the m^6^A abundance in aneuploidies did not follow the expression levels of transmethylases.

**Figure 2.**
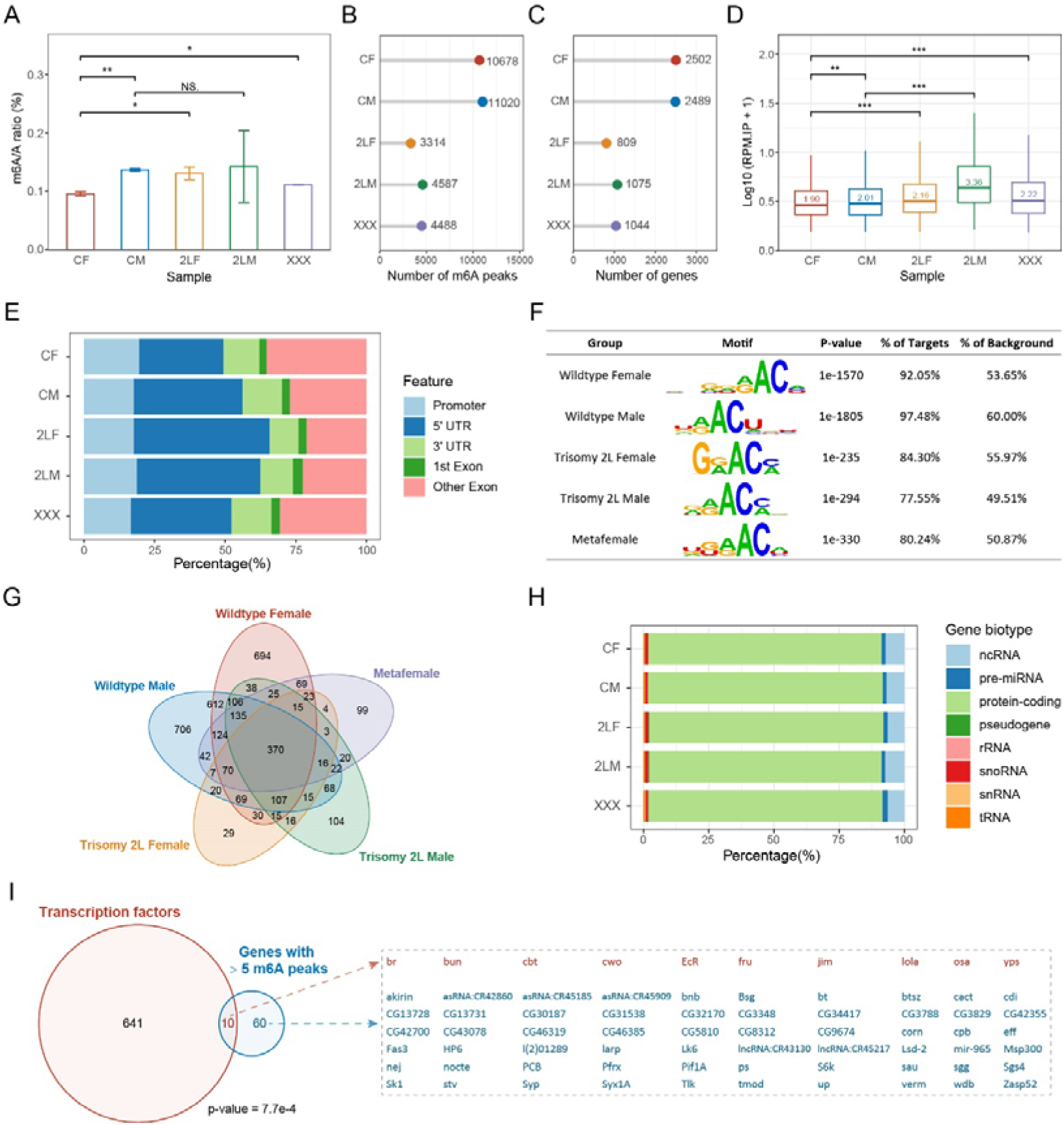
Overview of RNA m^6^A methylation in aneuploid Drosophila. (A) Global m^6^A abundance in third instar larvae of wildtype and aneuploid Drosophila. Data represent the mean of two independent experiments, each containing three or four biological replicates. Quantification was performed using EpiQuik m^6^A RNA Methylation Quantification Kit. Error bar indicates the standard error of the means (SEM). Student’s t test *p < 0.05, **p < 0.01. (B) The number of m^6^A peaks identified by MeRIP-Seq. (C) The number of m^6^A-modified genes obtained by annotating the peaks. (D) Expression levels of m^6^A modification sites in IP samples expressed as log10-transformed reads per million (RPM). The number on the boxplot indicates the median RPM of each sample. Mann-Whitney U test *p < 0.05, **p < 0.01, ***p < 0.001. (E) Percentages of peaks localized on different gene features. (F) The most enriched motifs obtained by de novo motif analysis of the m^6^A peaks. (G) Venn diagram showing the intersection of m^6^A-modified genes in each sample. (H) Gene biotypes of m^6^A-modified genes. (I) Venn diagram showing the intersection of transcription factors and m^6^A-modified genes with more than five m^6^A peaks in all samples. Seventy genes with more than five m^6^A peaks were listed on the right, with transcription factors in red and others in blue. P-value indicates one-tailed Fisher’s exact test. CF, wildtype female control; CM, wildtype male control; 2LF, trisomy 2L female; 2LM, trisomy 2L male; XXX, metafemale; 5[1UTR, 5[1 untranslated region; 3[1UTR, 3[1 untranslated region; MeRIP-Seq, m^6^A methylated RNA immunoprecipitation sequencing.

To obtain m^6^A mapping across the transcriptome of aneuploid Drosophila, whole larvae with different karyotypes were used for m^6^A methylated RNA immunoprecipitation sequencing (MeRIP-Seq). By identifying consistent peaks in two biological replicates, approximately ten thousand m^6^A peaks were found in wildtype females and males, whereas there were less than half the number of methylation sites in the three kinds of aneuploidies (Figure 2B). When these m^6^A sites were annotated to the genes, the changes in the number of m^6^A-marked genes in samples of different genotypes were in accordance with that of m^6^A peaks, in which there are about 2,500 m^6^A-marked genes in wildtype and about 1,000 m^6^A-modified genes in trisomies (Figure 2C). Both the number of m^6^A peaks and the number of m^6^A-modified genes are the least in trisomy 2L females (Figure 2B,C). The changes in the number of RNA methylation sites in aneuploids did not coincide with the changes of overall m^6^A abundance, but instead matched the expression of m^6^A components. We speculate that this may be caused by the nonuniformity and heterogeneity of RNA m^6^A modification, including the tissue specificity, the developmental specificity, the different numbers of m^6^A sites in one transcript, the different proportions of methylated transcripts, et cetera (Meyer et al., 2012; Meyer and Jaffrey, 2014; Zaccara et al., 2019). Counting the number of reads that mapped to m^6^A peaks in wildtype and aneuploidy MeRIP samples showed that the read levels at m^6^A sites in aneuploidies are significantly higher than that in wildtypes (Mann Whitney U test p-values < 0.001; Figure 2D); further analysis of the relative IP/Input ratios showed that the values of aneuploidies are still higher than wildtypes (Mann Whitney U test p-values < 0.001), which confirmed our hypothesis. These results suggest that RNA m^6^A modification exhibits greater heterogeneity in unbalanced genomes.

We next studied the overall characteristics of m^6^A methylation in aneuploid and control Drosophila. m^6^A peaks are localized along all autosomes and sex chromosomes, and the numbers of peaks are correlated with chromosome length (Figure 2—figure supplement 1A). Methylation sites are widely distributed on 5’UTR (30-50%), 3’UTR (10-15%), promoter (15-20%), and internal exons (20-35%), and the aneuploidies appeared to have a higher proportion of 5’UTR peaks and a lower proportion of exon peaks than the wildtype (Figure 2E). The highest values of density distributions of the length of exons containing m^6^A peaks are around 250 bp, and approximately 90% of the exons are longer than the typical 140 bp (Figure 2—figure supplement 1B). De novo motif analysis of m^6^A methylation peaks in each sample showed that the top-ranked motif is consistent with the conserved m^6^A motif DRACH (D=G/A/U, R=G/A, H=U/A/C; Figure 2F). In addition, enrichment analysis of the known motif DRACH showed that it is significantly enriched in all samples, and more than 97% of the m^6^A sites have this sequence (Figure 2—figure supplement 1C).

There are 370 common genes to which the m^6^A peaks were annotated in all genotypes (Figure 2G). We classified the m^6^A-modified genes in each sample and found that they were mostly protein-coding genes, whereas only about 7% were ncRNAs (Figure 2H). As described in previous studies (Meyer et al., 2012; Zaccara et al., 2019), most methylated RNAs have one m^6^A peak (Figure 2—figure supplement 1D-H). However, there are still some genes whose transcripts can be highly methylated and contain more than five m^6^A sites (Figure 2—figure supplement 1D-I). It was found that transcription factors were significantly enriched in 70 genes with more than 5 m^6^A peaks in both wildtype and aneuploidy (Fisher’s exact test p-value = 7.7e-4; Figure 2I; Figure 2—figure supplement 1I). Thus, specific types of genes may be preferentially targeted by m^6^A modification, such as transcriptional regulators (Kan et al., 2017). Analysis of the functions of m^6^A-modified genes in each sample revealed that they were enriched for a large number of common functions, among which the most significant were those related to morphogenesis, development, and growth (Figure 2—figure supplement 2A). The 141 GO terms shared by the five genotypes were summarized and found to include functions about metabolism, regulation, cellular processes, signaling pathways, behavior, and immune response (Figure 2—figure supplement 2B,C). Through pathway enrichment analysis of m^6^A-modified genes, 9 terms were found to be consistently enriched in all samples, including MAPK signaling pathway, Hippo signaling pathway, TGF-β pathway, etc. (Figure 2—figure supplement 2D-F).

### Differentially methylated peaks (DMPs) and their associated genes

We then searched for the differential methylation peaks (DMPs) in autosomal and sex chromosome aneuploidies compared with wildtype Drosophila of their corresponding sex. The number of DMPs and the changing trend of DMP methylation status are shown in the Figures (Figure 3A-D). In trisomy 2L females, the number of upregulated DMPs (1,131) is less than that of downregulated DMPs (1,602). In trisomy 2L males, the number of upregulated DMPs (1,521) is slightly higher than that of downregulated DMPs (1,259). In metafemales, the number of upregulated DMPs (1,393) is higher than that of downregulated DMPs (926) (Figure 3A-D). More DMP-associated genes were found in autosomal aneuploidies than in sex chromosome aneuploidy (trisomy 2L female = 2,148, trisomy 2L male = 2,055, metafemale = 1,771; Figure 3E). Also, the total number of peaks after the differential analysis would be different from the number of m^6^A peaks identified in the previous section, because the analysis software merged and recalculated the peaks (Stark and Brown, 2013).

**Figure 3.**
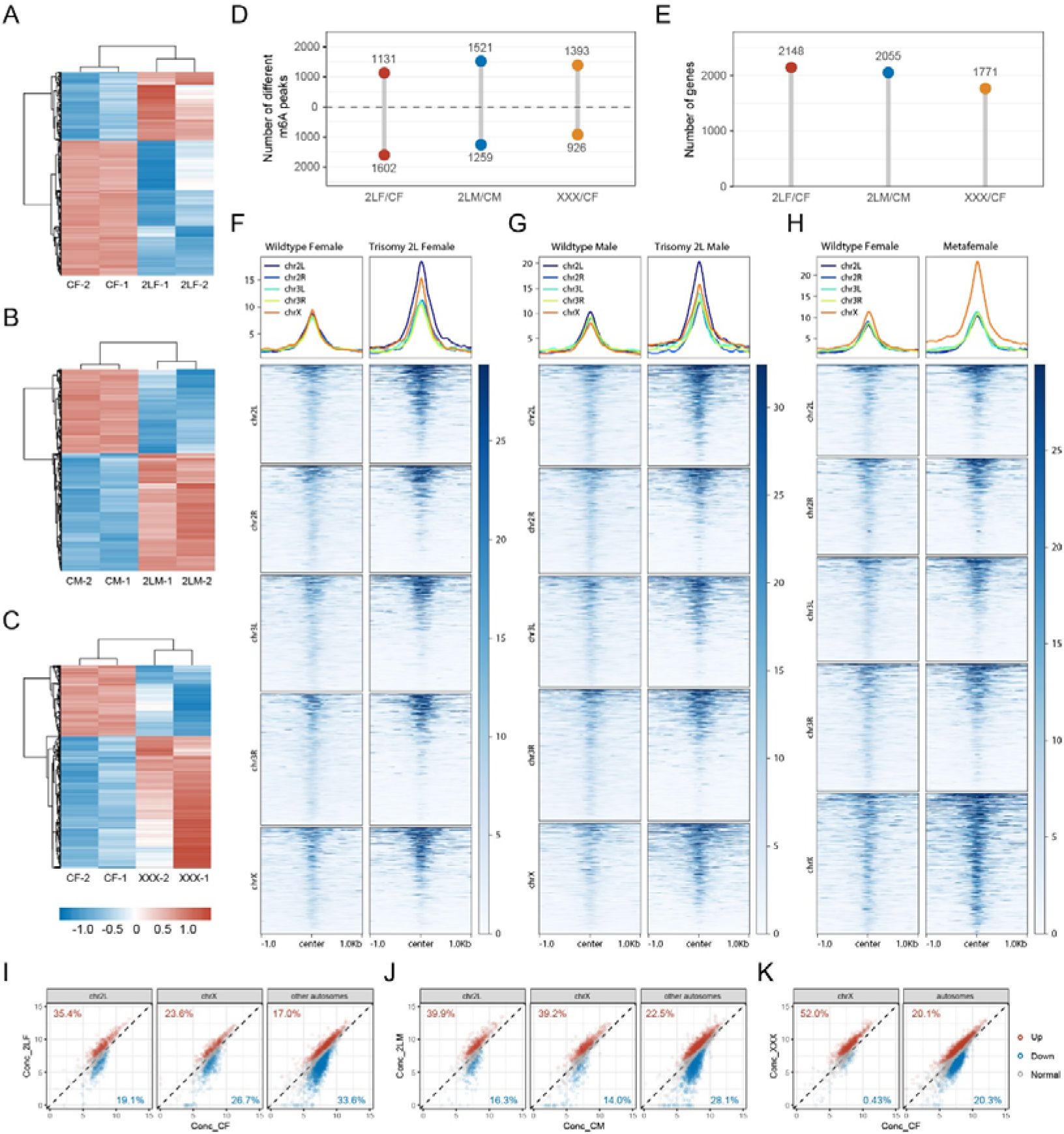
Differential m^6^A methylome analysis of aneuploid Drosophila. (A-C) Heatmaps of differentially methylated peaks (DMPs) in trisomy 2L females (A), trisomy 2L males (B) and metafemales (C), and their corresponding control groups. The threshold of significance was p-value ≤ 0.1. (D) The number of DMPs. The threshold of significance was set to p-value ≤ 0.1. The numbers above the horizontal dashed lines indicate peaks with upregulated methylation levels, and the numbers below indicate peaks with downregulated methylation levels. (E) The number of DMP-associated genes. (F-H) Profiles and heatmaps illustrating the density of m^6^A-modified reads at the DMP positions in trisomy 2L females (F), trisomy 2L males (G), metafemales (H), and their corresponding controls. The DMPs were divided into five groups according to the chromosomes they located. (I-K) Scatter plots showing the concentration of reads at methylation sites on different chromosomes in trisomy 2L females (I), trisomy 2L males (J), and metafemales (K). Red points indicate significantly up-regulated m^6^A peaks, blue points indicate significantly down-regulated m^6^A peaks, and gray points indicate m^6^A peaks without significant changes. The percentages of DMPs on cis and trans chromosomes were indicated in the corners of the plots. CF, wildtype female control; CM, wildtype male control; 2LF, trisomy 2L female; 2LM, trisomy 2L male; XXX, metafemale.

DMPs are distributed on all chromosomes (Figure 3—figure supplement 1A). Specifically, the MeRIP-Seq signal intensity at the location of DMPs showed that the m^6^A-marked reads in trisomies are denser than those of wildtypes, especially on the triple chromosomes (Figure 3F-H). The density of reads on chromosome 2L of trisomy 2L Drosophila is higher than that on other chromosomes, while the density of reads on chromosome X of metafemales is significantly higher than that of autosomes (Figure 3F-H). Previous studies on aneuploid Drosophila showed that the expression levels of genes on chromosomes with increased copy numbers were mostly similar to those in diploids, and the expression of genes on unvaried chromosomes were widely inverse regulated (Sun et al., 2013c; Birchler, 2016; Zhang et al., 2023). It can be concluded that the changes in the density of reads at DMP sites are not due to a direct gene dosage effect. Similar results were obtained by counting the methylation status of cis and trans transcripts (Figure 3I-K). Overall, the transcripts of genes located on the varied chromosomes in three trisomies have more up-regulated DMPs than down-regulated DMPs, which is more obvious in metafemales (Figure 3I-K). In trisomy 2L females and males, genes localized on other autosomes have a higher proportion of down-regulated DMPs, while the X-linked trans genes show sexual dimorphism and X chromosome-specific response to genomic imbalance. (Figure 3I,J). For metafemales, the transcripts of trans genes have similar numbers of up-regulated and down-regulated DMPs (Figure 3K). Therefore, cis genes in trisomy generally possessed a higher proportion of up-regulated DMP-associated transcripts, whereas the methylation states of trans genes are variable, depending on the identities of varied chromosomes and their genomic locations. The above results give us reason to speculate that the changes of m^6^A methylation may affect dosage compensation of cis-genes and inverse dosage modulation of trans-genes in aneuploidy.

The distribution of DMPs along gene features includes 5’UTR (30-35%), 3’UTR (20-25%), promoter (10-25%), and internal exon (25-35%), among which the proportion of 3’UTR is higher than that of all m^6^A sites (Figure 3—figure supplement 1B). The length distributions of exons with DMPs are similar to that of all m^6^A-modified exons (Figure 3—figure supplement 1C). De novo motif analysis of the sequences where DMPs are located or enrichment analysis of the known motif DRACH both showed that DMP sites are enriched for the conserved motif of m^6^A (Figure 3—figure supplement 1D,E). There are also similarities between the types of DMPs-associated genes and m^6^A-marked genes (Figure 3—figure supplement 1F). The DMP-associated genes of the three aneuploidies have complex overlapping relationships (Figure 3—figure supplement 1G). Most DMP-associated genes have one differentially methylated site, and only a few genes contained more than three DMPs (Figure 3—figure supplement 1H-J). There are 1,042 common DMP-associated genes in all trisomies, of which 71 were transcription factors and were significantly enriched (Fisher’s exact test p-value = 6.9e-5; Figure 4A,B). We speculate that these differentially methylated transcription factors may play an important role in the regulation of gene expression and development of aneuploidy.

**Figure 4.**
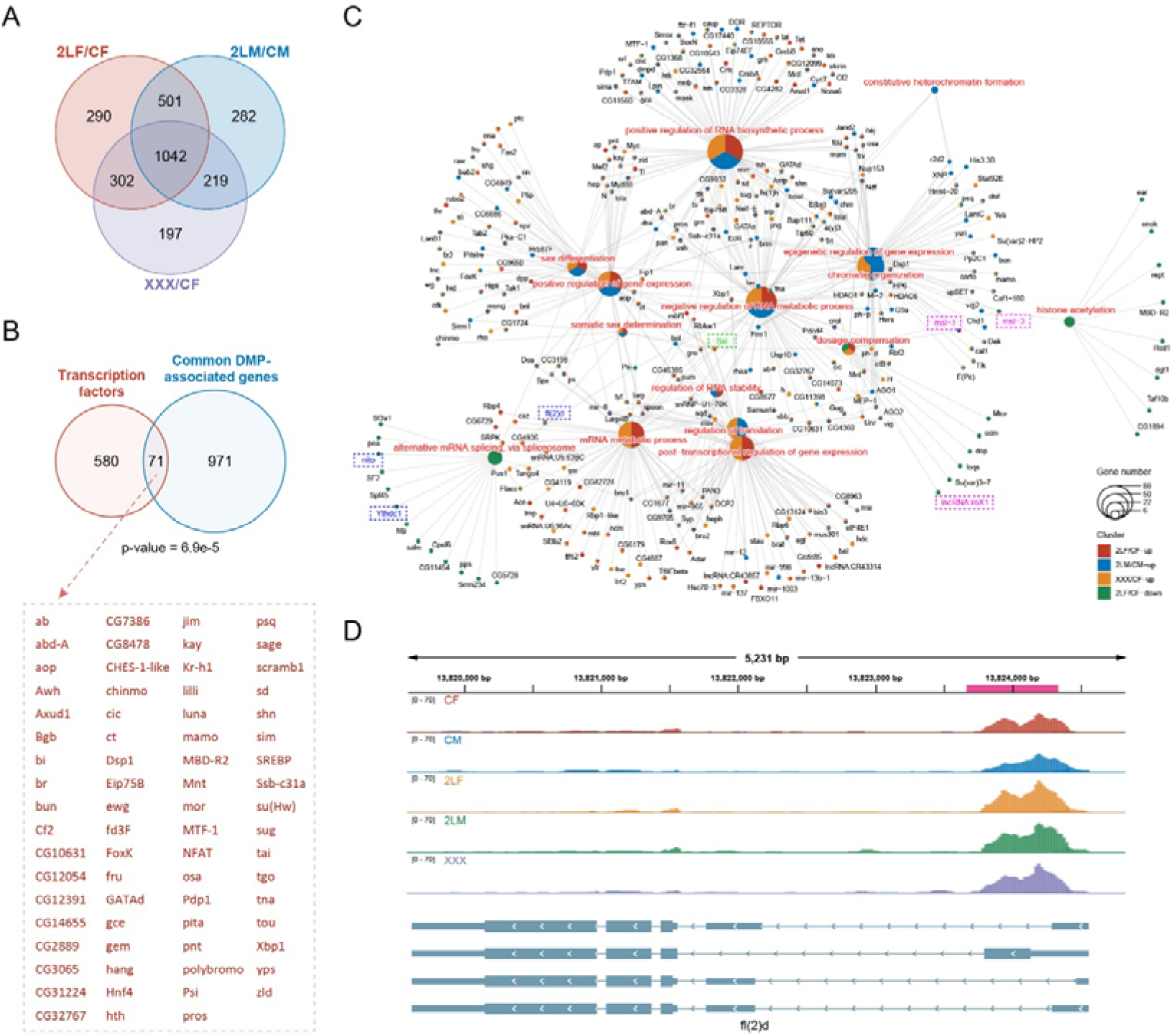
Differentially methylated peak (DMP) associated genes and their functions. (A) Venn diagram showing the number of common DMP-associated genes in three types of aneuploidies compared with wildtypes. (B) Venn diagram showing the intersection of transcription factors and the common DMP-associated genes in all comparisons. Transcription factors with DMPs were listed on the right. P-value indicates one-tailed Fisher’s exact test. (C) Network showing the functions related to expression regulation and dosage compensation enriched by DMP-associated genes. The color of the nodes indicates the comparison, and the size of the function nodes represents the number of DMP-associated genes connected with them. (D) Genome browser example of fl(2)d for indicated MeRIP-seq data. Steelblue color represents input reads, while other colors represent IP reads. Signals were displayed as the mean CPM of two biological replicates. The gene architectures were shown at the bottom. The magenta rectangles at above represent differentially methylated peak. CF, wildtype female control; CM, wildtype male control; 2LF, trisomy 2L female; 2LM, trisomy 2L male; XXX, metafemale.

Functional enrichment analysis showed that genes with up-regulated DMPs in three trisomies were enriched for 153 common functions, while the genes with down-regulated DMPs shared only 6 functions (Figure 4—figure supplement 1A,B). Similarly, genes with up-regulated DMPs were enriched for more consistent pathways in all aneuploids (Figure 4—figure supplement 1C,D). Most of the functions of DMP-associated genes are the same as those of m^6^A-marked genes, and these genes are also enriched for cell fate commitment, dorsal/ventral pattern formation, sex differentiation, post-transcriptional regulation of gene expression, and other additional GO terms. We found that many DMP-associated genes have functions related to gene expression regulation and dosage compensation, including some components of MSL complex (msl-1, msl-3, lncRNA:roX1) and the major sex-determining factor Sxl (Figure 4C). We also noticed that m^6^A component genes fl(2)d, nito, and Ythdc1 themselves are differentially methylated in aneuploidy (Figure 4C), for example, there was a common significantly upregulated m^6^A peak in the 5’UTR of the transcripts of fl(2)d in three aneuploidies (Figure 4D). These data suggest that m^6^A dynamics in aneuploids may be involved in the expression regulation of unbalanced genomes through various genes with regulatory functions.

### Relationships between m A and dosage-related modulation of gene expression in aneuploidy

To explore the relationships between RNA m^6^A modification and gene expression regulation under genomic imbalance, we combined the results of MeRIP-Seq and RNA-seq, and performed a comprehensive analysis. The results showed that about 12%-15% of genes differentially expressed in aneuploid Drosophila also belong to DMPs-associated genes (Figure 5A-C). However, except for trisomy 2L males, the DEGs of the other two trisomies did not show enrichment of differential m^6^A methylation (Fisher’s exact test p-values: trisomy 2L male = 0.012, trisomy 2L female and metafemale > 0.05). There are 55 genes that are both differentially expressed and differentially methylated in all aneuploidies (Figure 5D). By analyzing the functions of genes that are simultaneously differentially expressed and differentially methylated, we obtained a number of consistent biological functions, including post-embryonic animal morphogenesis, cell communication, signal transduction, response to stimulation, and so on (Figure 5E), possibly reflecting the important roles of m^6^A-regulated expression in cell signaling and development of organisms.

**Figure 5.**
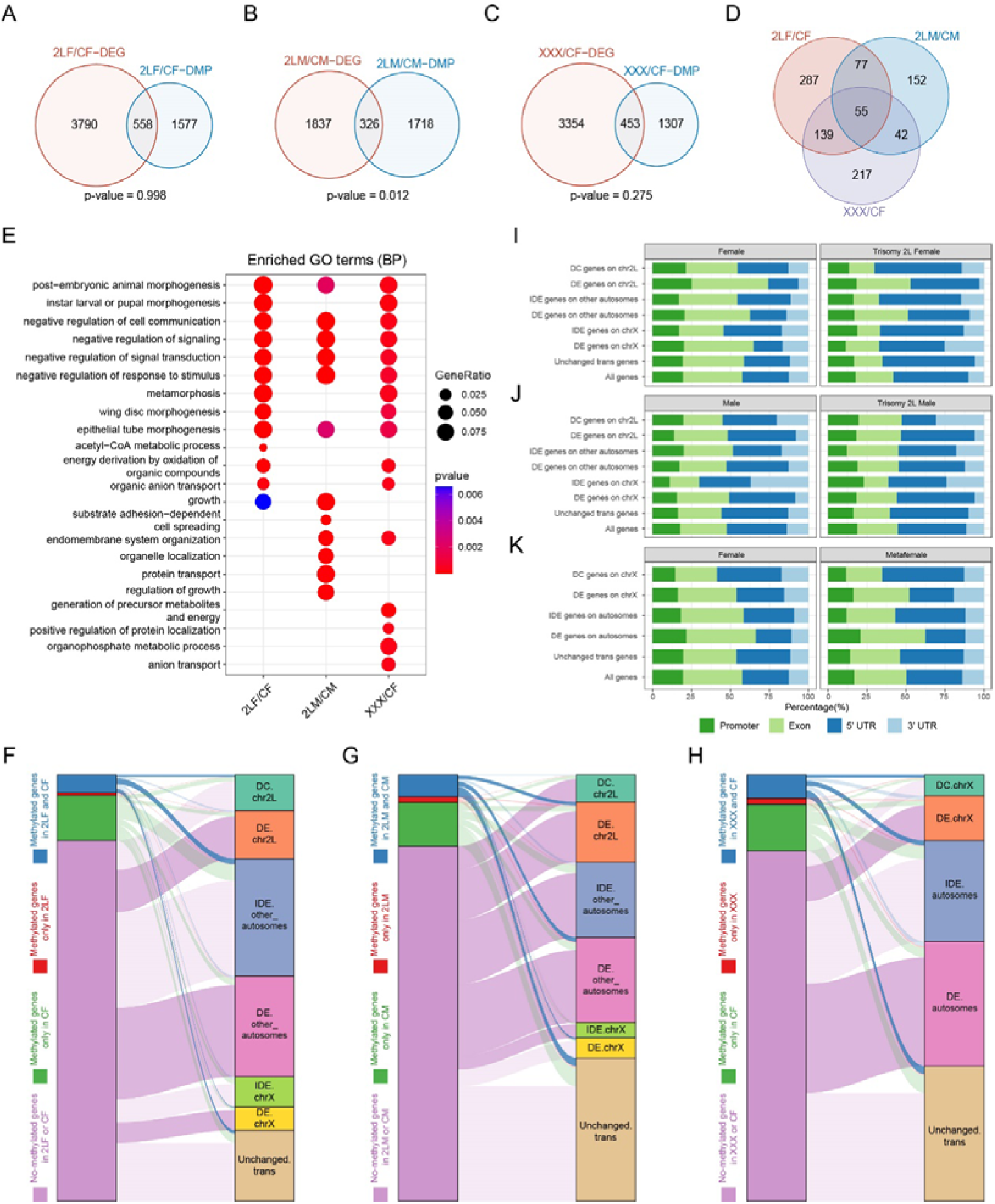
Relationships between RNA m^6^A methylation and gene expression in aneuploidy. (A-C) Venn diagrams showing the intersections of differentially expressed genes (DEGs) and differentially methylated peak (DMP) associated genes in trisomy 2L females (A), trisomy 2L males (B), and metafemales (C) compared with their corresponding controls. P-values indicate one-tailed Fisher’s exact tests. (D) The common differentially expressed and differentially methylated genes in all groups. (E) Functional enrichment analysis of simultaneously differentially expressed and differentially methylated genes. Top 10 enriched GO terms (Biological Process) with p-value < 0.1 in each comparison were shown. (F-H) Sankey diagrams showing the relationships between genes with different m^6^A-modified states and genes with canonical dosage effect (DE), dosage compensation (DC), and inverse dosage effect (IDE) in trisomy 2L females (F), trisomy 2L males (G), and metafemales (H). Enrichment analysis was performed on each two groups of genes, and deep color lines indicate significant connection relationships (Fisher’s exact test p-value < 0.05). (I-K) Gene feature distributions for m^6^A peaks on genes with canonical DE, DC, and IDE in trisomy 2L females (I), trisomy 2L males (J), metafemales (K), and their corresponding controls. CF, wildtype female control; CM, wildtype male control; 2LF, trisomy 2L female; 2LM, trisomy 2L male; XXX, metafemale; DEG, differentially expressed gene; DMP, differentially methylated peak; DE, dosage effect; DC, dosage compensation; IDE, inverse dosage effect. Canonical DE refers to ratio > 1.25, DC stands for 0.8 < ratio < 1.25, IDE stands for 0.5 < ratio < 0.8, and unchanged refers to 0.8 < ratio < 1.25.

By analyzing the crossover of different groups of m^6^A-modified genes and genes with canonical dosage effect, dosage compensation, and inverse dosage effect, we further revealed the relationships between m^6^A-modified genes and dosage-related effects in aneuploidy (Figure 5F-H). The results showed that in trisomy 2L females, cis dosage compensation genes, trans autosomal dosage effect genes, and trans unchanged genes are significantly enriched in m^6^A group with methylation in both trisomy and control; meanwhile, cis dosage effect genes, other autosomal dosage effect genes, and X-linked dosage effect genes are enriched in the group without methylation in both trisomy and control (Figure 5F). On the contrary, in trisomy 2L males, cis dosage effect genes, trans dosage effect genes, and trans unchanged genes are significantly enriched in m^6^A group whose genes are methylated in both trisomy and control; while the group of genes that are not methylated at all is enriched for cis dosage effect genes, cis dosage compensation genes, trans inverse dosage effect genes, and other autosomal dosage effect genes (Figure 5G). Metafemales performed similarly to trisomy 2L female, with significant enrichment of cis dosage compensation genes, autosomal trans dosage effect genes, and trans unchanged genes in m^6^A group where both trisomy and control genes are methylated; and the group without methylation is enriched for cis dosage effect genes and trans dosage effect genes (Figure 5H). These results indicated that m^6^A-modified genes in aneuploid Drosophila females are mainly related to dosage compensation and inverse dosage effect, while genes not modified by m^6^A are mainly related to direct dosage effect. Male aneuploids did not follow this trend.

Furthermore, the distributions of m^6^A modification sites along gene features in genes with classical dosage-related effects were also studied (Figure 5I-K). We found that in wildtype females, trisomy 2L females, and metafemales, a higher proportion of m^6^A sites on genes with canonical dosage compensation and inverse dosage effect are distributed in the 5’UTR than genes with dosage effects (Figure 5I,K). At the same time, this proportion is higher in trisomy than in wildtype. In contrast to females, trisomy 2L males have a higher proportion of 5’UTR RNA m^6^A modification on dosage effect genes (Figure 5J). The above study again demonstrates the sexual dimorphism of RNA m^6^A modification in response to aneuploidy. In addition, it has been suggested that m^6^A residues in the 5’UTR region may have unique regulatory functions (Luo et al., 2014; Meyer and Jaffrey, 2014). Therefore, we hypothesized that the high level of 5’UTR m^6^A modification on genes with dosage compensation and inverse dosage effect might regulate the down-regulation of these genes.

Dozens of dosage-sensitive modifiers have been identified in Drosophila, whose dosage changes can negatively or positively regulate the gene expression across the whole genome like aneuploidy (Birchler et al., 2001; Birchler and Veitia, 2007). Protein-protein interaction (PPI) networks between dosage-sensitive modifiers and differentially expressed genes (DEGs) were constructed to investigate whether RNA m^6^A modification could participate in the gene regulatory networks in unbalanced genomes through these regulators (Figure 5—figure supplement 1A-C). In the three types of aneuploidies, there are more dosage-sensitive modifiers with up-regulated DMPs than with down-regulated DMPs. The regulators ox and Vha55 without DMPs interact with a large number of significantly up-regulated DEGs in trisomy 2L females and metafemales, whereas these two regulators interact with a small number of down-regulated DEGs in trisomy 2L males. Up-regulated DMP-associated regulators wg, Uba1, ap, Atg1, osa, rdx, and sd in trisomy 2L females mainly interacted with significantly down-regulated DEGs (Figure 5—figure supplement 1A); and up-regulated DMP-associated regulators Uba1, osa, and sd in metafemales also interacted with down-regulated DEGs (Figure 5—figure supplement 1C). Up-regulated DMP-associated regulators wg, Kr-h1, and Trl mainly interacted with up-regulated DEGs in trisomy 2L males (Figure 5—figure supplement 1B). In addition, a preponderance of significantly down-regulated DEGs interacted with dosage-sensitive modifiers is observed on trans chromosomes of all Drosophila trisomies (Figure 5—figure supplement 1A-C).

### Alterations of m A may be involved in differential alternative splicing in imbalanced genomes

Next, we analyzed the differential alternative splicing events in aneuploid Drosophila. More than 1,000 differential splicing events have been identified in different aneuploids, and about one-third of them are of the type of skipped exon (SE) (Figure 6—figure supplement 1A-C). The biological functions of differentially spliced transcripts are mainly involved in macromolecular fiber organization, locomotion, and growth (Figure 6—figure supplement 1D), and these genes are enriched in heterogeneous pathways in different aneuploidies (Figure 6—figure supplement 1E). Notably, we found that genes with DMPs are significantly enriched for differential alternative splicing events in three kinds of aneuploid Drosophila (Fisher’s exact test p-values < 0.05; Figure 6A-C). Among the genes whose transcripts are differentially spliced, 27%-32% are also differentially methylated (Figure 6A-C). There are 67 genes with both differential alternative splicing and differential m^6^A methylation in all aneuploidies (Figure 6D). The functions of these genes are similar to those of all differentially spliced genes (Figure 6E), but more consistent KEGG pathways are enriched, including endocytosis, mTOR signaling pathway, Hedgehog signaling pathway, and so on (Figure 6F).

**Figure 6.**
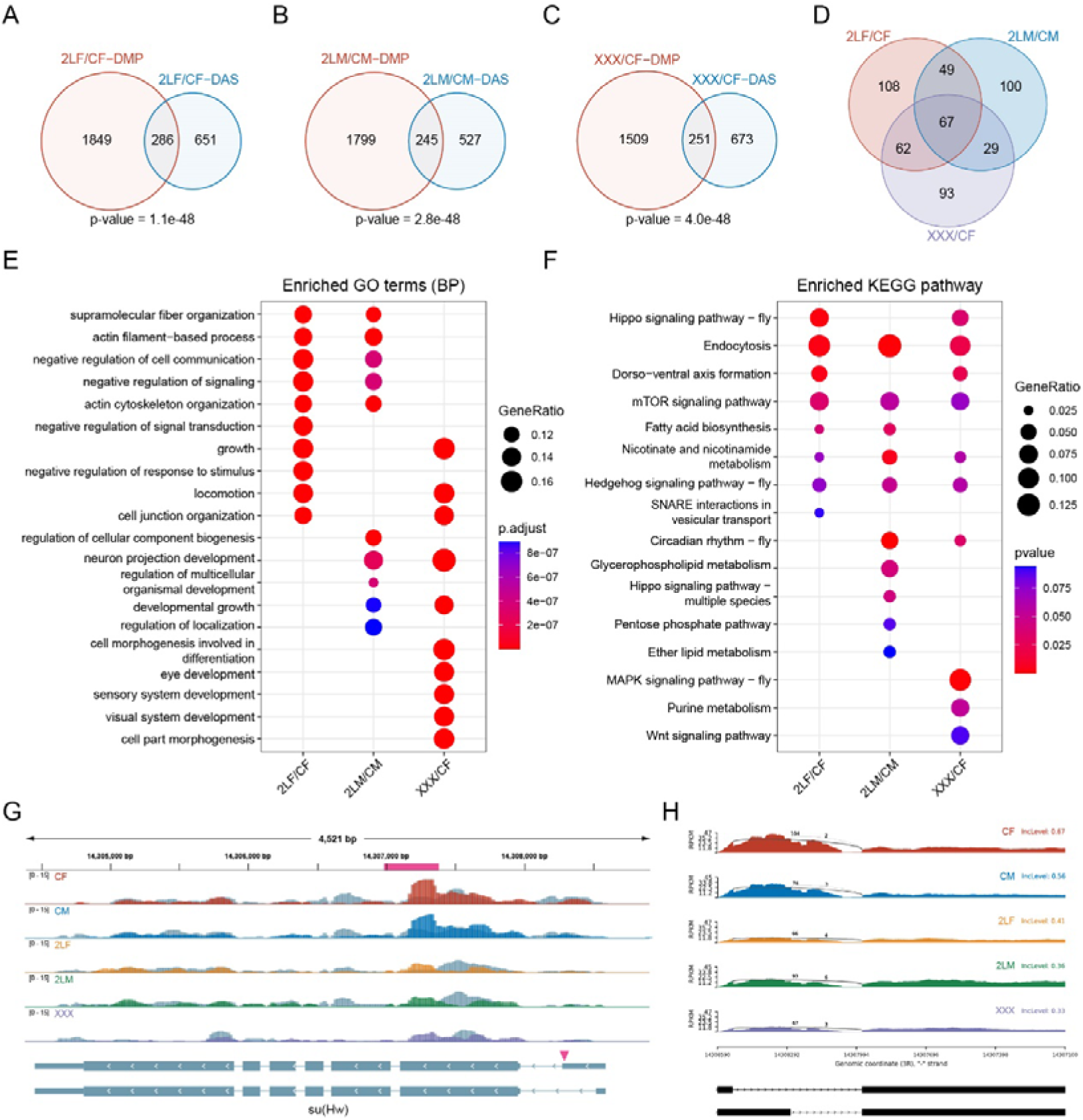
Combined analysis of differential alternative splicing and differential methylation. (A-C) Venn diagrams showing the intersections of differential alternative splicing (DAS) genes and differentially methylated peak (DMP) associated genes in trisomy 2L females (A), trisomy 2L males (B), and metafemales (C) compared with their corresponding controls. P-values indicate one-tailed Fisher’s exact tests. (D) The common differentially alternatively spliced and differentially methylated genes in all groups. (E) Functional enrichment analysis of simultaneously differentially alternatively spliced and differentially methylated genes. Top 10 enriched GO terms (Biological Process) with p-value < 0.05 in each comparison were shown. (F) KEGG pathway enrichment analysis of simultaneously differentially alternatively spliced and differentially methylated genes. Top 10 enriched pathways with p-value < 0.1 in each comparison were shown. (G) Genome browser example of su(Hw) for indicated MeRIP-Seq data. Steelblue color represents input reads, while other colors represent IP reads. Signals were displayed as the mean CPM of two biological replicates. The gene architecture was shown at the bottom (only two representative transcript isoforms were shown). The magenta rectangle at above represents DMP. The magenta arrowhead indicates the position of differential alternative splicing. (H) Sashimi plot depicting RNA sequencing reads and exon junction reads at the position where the differential splicing events occurs on su(Hw). The gene model was shown below. One of the biological replicates was chosen for representation. CF, wildtype female control; CM, wildtype male control; 2LF, trisomy 2L female; 2LM, trisomy 2L male; XXX, metafemale; DAS, differential alternative splicing; DMP, differentially methylated peak; CPM, Counts per million; RPKM, Reads per kilobase per million mapped reads.

In all trisomy Drosophila, 10 genes are shown to be differentially expressed, with their transcripts also being differentially spliced and differentially methylated. Three of them are illustrated below [su(Hw), Ppn, and CG13124]. su(Hw) is a component of the gypsy chromatin insulator complex, which is a regulatory element that establish independent domains of transcriptional activity (Roseman et al., 1993). Due to the close relationship between su(Hw) and second-site modifiers (Rabinow and Birchler, 1989), and BEAF-32, which is also an insulator DNA-binding protein, has been proposed as a possible inverse dosage regulator (Gurudatta et al., 2012; Zhang et al., 2021b), We speculated that the transcription factor su(Hw) may also be a dosage-sensitive regulator. The data showed that the transcription levels of su(Hw) are up-regulated in all three aneuploidies, and there is a consistent m^6^A modification site with significantly down-regulated methylation (Figure 6G). At the same time, the transcripts of this gene have a common alternative 5’ splice site (A5SS), and its inclusion levels in trisomies are down-regulated, that is, more short transcript isoforms are generated (Figure 6G). Ppn gene encodes an essential extracellular matrix protein that influences cell rearrangements. The expression level of Ppn and its m^6^A methylation in 5’UTR region are both significantly up-regulated in trisomies (Figure 6—figure supplement 1F). In addition, an alternatively spliced exon is significantly less frequently skipped in all aneuploid Drosophila (Figure 6—figure supplement 1G). The third gene, CG13124, which may be involved in involved in regulation of translational initiation, is up-regulated in aneuploids. Two m^6^A sites in its 5’UTR region are methylated at higher levels in all aneuploid Drosophila (Figure 6—figure supplement 1H). Its transcripts also have a common significantly different exon-skipping event (Figure 6—figure supplement 1I).

These results indicate that there are complicated relationships among RNA m^6^A modification, gene expression, and alternative splicing under the condition of genome imbalance. RNA splicing seems to be more closely related to m^6^A methylation than gene transcription. The m^6^A sites located in 5’UTR show remarkable changes in methylation levels in aneuploid Drosophila, and may be involved in the regulation of some differential alternative splicing events, such as exon skipping.

### Interactions between m A and Drosophila Male Specific Lethal (MSL) complex

Previous studies have shown that m^6^A components are involved in regulating the alternative splicing of sex-determining gene Sxl in Drosophila, and the deficiency of m^6^A writers or readers will lead to the reduction of female-specific isoforms of Sxl (Haussmann et al., 2016; Lence et al., 2016; Kan et al., 2017). Sxl is also a direct target of RNA m^6^A modification (Kan et al., 2017). We found that Sxl transcripts are both differentially methylated and differentially spliced in three kinds of aneuploid Drosophila (Figure 7A,B). For trisomy 2L females and metafemales, two common m^6^A peaks are significantly up-regulated in the 5’UTR region of Sxl. However, trisomy 2L males have a significantly down-regulated m^6^A peak in the 5’UTR (Figure 7A). Meanwhile, multiple junctions of Sxl transcripts undergo complicated alternative splicing in aneuploid Drosophila, including SE, A5SS, alternative 3’ splice site (A3SS), and mutually exclusive exons (MXE) types (Figure 7A). By checking the distributions of RNA sequencing reads near the male-specific exon (namely the third exon), it can be observed that there are almost no mapped reads on the third exon in wildtype females, trisomy 2L females, and metafemales, while the reads mapped to the second and fourth exons are highly prevalent (Figure 7B). On the contrary, wildtype males and trisomy 2L males have a substantial number of reads on the third exon, accompanied by a smaller number of reads on the adjacent two exons (Figure 7B). Notably, we found that a small number of RNA-seq reads aligned to the male-specific exon appeared in trisomy 2L females, which was identified as an SE-type differential alternative splicing event (FDR = 7.4e-7; Figure 7B). This variation may be related to the abnormal m^6^A methylation levels under the condition of genomic imbalance.

**Figure 7.**
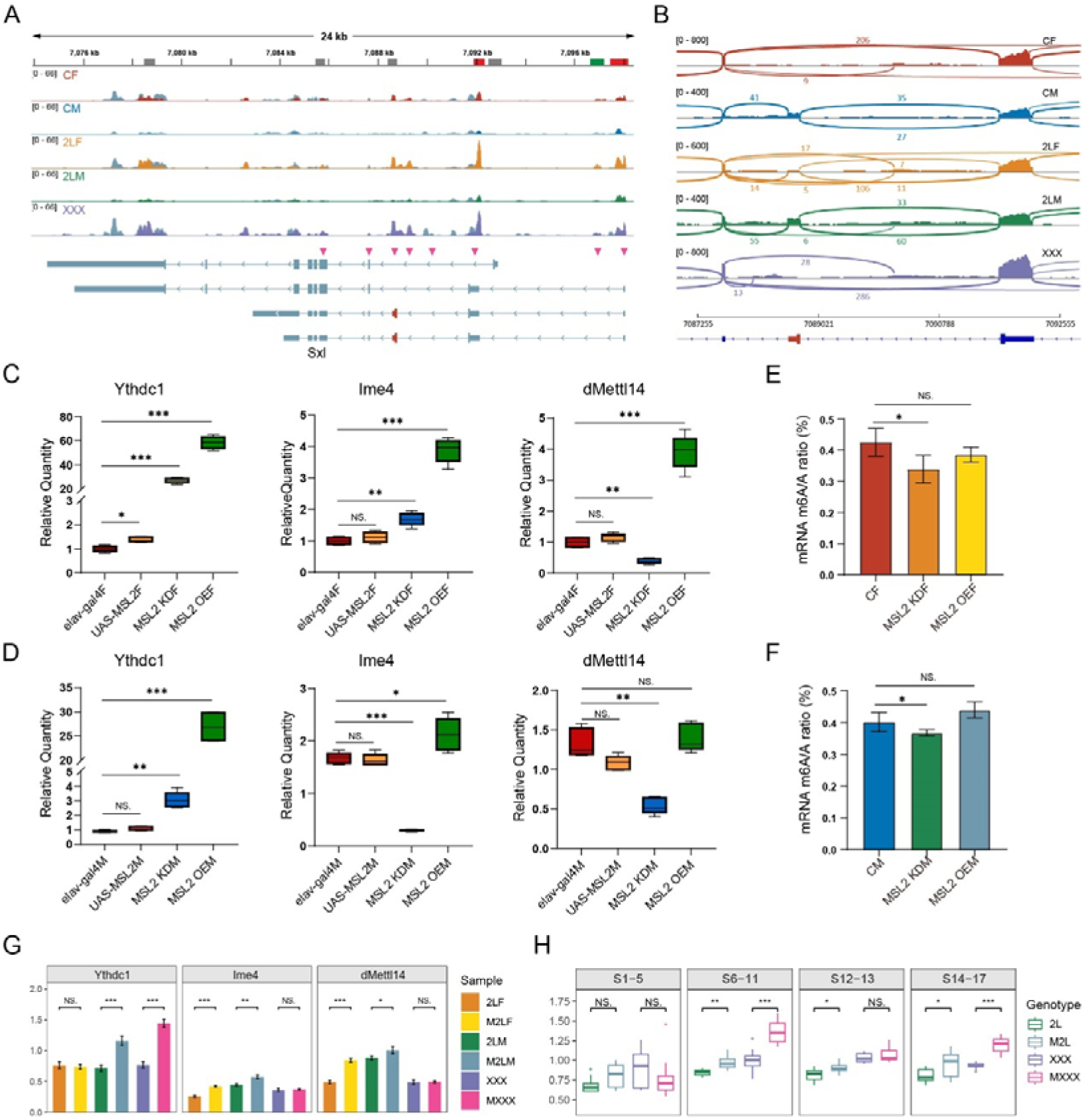
Interactions between m^6^A and Drosophila Male Specific Lethal (MSL) complex. (A) Genome browser example of Sxl for indicated MeRIP-Seq data. Steelblue color represents input reads, while other colors represent IP reads. Signals were displayed as the mean CPM of two biological replicates. The gene architecture was shown at the bottom (only four representative transcript isoforms were shown). The rectangles at above represent m^6^A peaks, where red indicates up-regulated DMPs, green indicates down-regulated DMP, and gray indicates no significant changes. The magenta arrowheads indicate the positions of alternative splicing. (B) Sashimi plot depicting RNA sequencing reads and exon junction reads at the position where the differential splicing events occurs on Sxl. The gene model was shown below, with the third exon indicated in red. One of the biological replicates was chosen for representation. CF, wildtype female control; CM, wildtype male control; 2LF, trisomy 2L female; 2LM, trisomy 2L male; XXX, metafemale. (C,D) RT-qPCR analysis of mRNA levels of m^6^A components in the heads of MSL2 transgenic female (C) and male (D) Drosophila adults. (E,F) Abundance of mRNA m^6^A modification in the heads of MSL2 transgenic females (E) and males (F). Student’s t test *p < 0.05, **p < 0.01, ***p < 0.001. MSL2 KDF, MSL2 neural-knockdown female; MSL2 KDM, MSL2 neural-knockdown male; MSL2 OEF, MSL2-overexpressed female; MSL2 OEM, MSL2-overexpressed male. (G) RT-qPCR analysis of mRNA levels of m^6^A regulators in the brains of trisomy and MSL2-overexpressed trisomy Drosophila larvae. Student’s t test *p < 0.05, **p < 0.01, ***p < 0.001. 2LF, trisomy 2L female; 2LM, trisomy 2L male; XXX, metafemale; M2LF, MSL2-overexpressed trisomy 2L female; M2LM, MSL2-overexpressed trisomy 2L male; MXXX, MSL2-overexpressed metafemale. (H) The expression levels of Ime4 in trisomy and MSL2-overexpressed trisomy embryos represented by relative fluorescence intensity of probes. The expression of wildtype embryos was set as one. Sample size = 10. Student’s t test *p < 0.05, **p < 0.01, ***p < 0.001. 2L, trisomy 2L; M2L, MSL2-overexpressed trisomy 2L; XXX, metafemale; MXXX, MSL2-overexpressed metafemale.

The abnormal expression of m^6^A components reduces the survival of female Drosophila, which is thought to be probably caused by the expression of downstream MSL complex (Haussmann et al., 2016). To investigate the interplay between the MSL complex and m^6^A modification, we examined the responses of m^6^A regulators in transgenic Drosophila strains with MSL2 mutation or overexpression (Figure 7—figure supplement 1A-E; Figure 7C,D). The results revealed significant changes in the expression profiles of m^6^A regulators in MSL2 transgenic strains, especially the m^6^A reader protein Ythdc1, which increased tens of fold in MSL2 knockdown and overexpressed Drosophila (Figure 7C,D; Figure 7—figure supplement 1E). We also observed that the trends of Ime4 expression in females and males were the same in MSL2-overexpressed Drosophila, whereas there was obvious sexual dimorphism in MSL2-knockdown samples (Figure 7C,D), which may be due to the ectopic assembly of MSL complex in MSL2-overexpressed females (Zhang et al., 2021a). In addition, we also examined the expression levels of MSL2 when Ythdc1 was knocked down, and found that MSL2 was also significantly increased in females (Figure 7—figure supplement 1F). All these results strongly suggest a potential relationship between MSL2 and Ythdc1. Next, we further compared the overall abundance of m^6^A on mRNA in MSL2 transgenic and wildtype Drosophila (Figure 7E,F).

The results showed that mRNA m^6^A levels were significantly decreased with MSL2 knockdown in females and males, but overexpression of MSL2 failed to exert a discernible effect on m^6^A abundance (Fig.7E,F).

In the next, we investigated the expression of m^6^A regulators in aneuploid Drosophila overexpressing MSL2, according the results that the MSL complex could be regulated directly or indirectly by m^6^A modification, and the sexual dimorphism of RNA m^6^A modification in response to aneuploidy. It is found that the transcription levels of m^6^A regulators are significantly up-regulated in the brains of most aneuploid larvae that overexpressed MSL2 (Figure 7G; Figure 7—figure supplement 1I). These results are not completely consistent with the quantitative results when MSL2 was overexpressed in diploids, which may be related to the effect of unbalanced genomes. We also used TSA-FISH to detect the expression and distribution of m^6^A components during embryogenesis in aneuploidies overexpressing MSL2 (Figure 7H; Figure 1—figure supplement 2). The subembryonic and subcellular distributions of mRNAs for m^6^A methyltransferases and reading protein did not appear to be affected by ectopic expression of MSL2 in aneuploid Drosophila embryos (Figure 1—figure supplement 2A-F; Figure 1—figure supplement 2A’-F’). But the relative expression of m^6^A components showed diverse dynamics, among which the levels of the most important methyltransferase Ime4 are significantly up-regulated in autosomal trisomy and sex chromosome trisomy with MSL2 overexpression (Figure 7H; Figure 1—figure supplement 2A”-F”). Considering that unbalanced genomes can affect the expression of MSL complex subunits, and the above results show that there is a close relationship between MSL complex and m^6^A modification, we speculate that unbalanced genomes may influence the expression of m^6^A regulators, possibly through the MSL complex.

### Relationship of H4K16Ac with m A modification

As an important component of the MSL complex, MOF is a histone acetyltransferase that specifically acetylates histone H4 at lysine 16 (H4K16Ac), thereby affecting chromatin structure and functions, and activating transcription (Kind et al., 2008; Conrad et al., 2012). Previous studies have elucidated the regulatory role of m^6^A in histone modifications, and histone 3 lysine 36 trimethylation (H3K36me3) also plays a role in recruiting methyltransferase complexes to deposit m^6^A markers on RNA (Wang et al., 2018; Huang et al., 2019; Li et al., 2020). To investigate the functions of MOF-mediated H4K16Ac on RNA m^6^A modification, we analyzed RNA-seq data from Drosophila strains overexpressing MOF (Figure 8A; Figure 7—figure supplement 1J-L). We found obvious changes in the expression of m^6^A regulators in strains overexpressing MOF, with the levels of almost all m^6^A regulators being elevated (Figure 8A). It was also observed that after overexpression of MOF, the increasing trends of m^6^A regulators in females and males were not exactly the same, mostly showing more pronounced increases in females, except for Mettl3 (Ime4) and dMettl14. According to the results of RT-qPCR, the expression of Ythdc1 was decreased in MOF knockdown strains and increased in MOF overexpression strains (Figure 8B; Figure 7—figure supplement 1J); meanwhile, the expression levels of MOF were significantly reduced in Ythdc1 neural-knockdown strains (Figure 8C). These results were further verified by polytene chromosome immunofluorescence experiments (Figure 8—figure supplement 1E, F). In males, the expression level of MOF was significantly decreased in Ythdc1 neural-knockdown Drosophila; while in Ythdc1 neural-knockdown females, the expression level of MOF was not significantly changed compared with wildtype females (Figure 8—figure supplement 1E, F). These results suggest that MOF-mediated acetylation modification can have a certain effect on RNA m^6^A methylation in a sexually dimorphic manner, which may be due to the endogenous MSL complex and the imbalance of X chromosome dosage in males.

**Figure 8.**
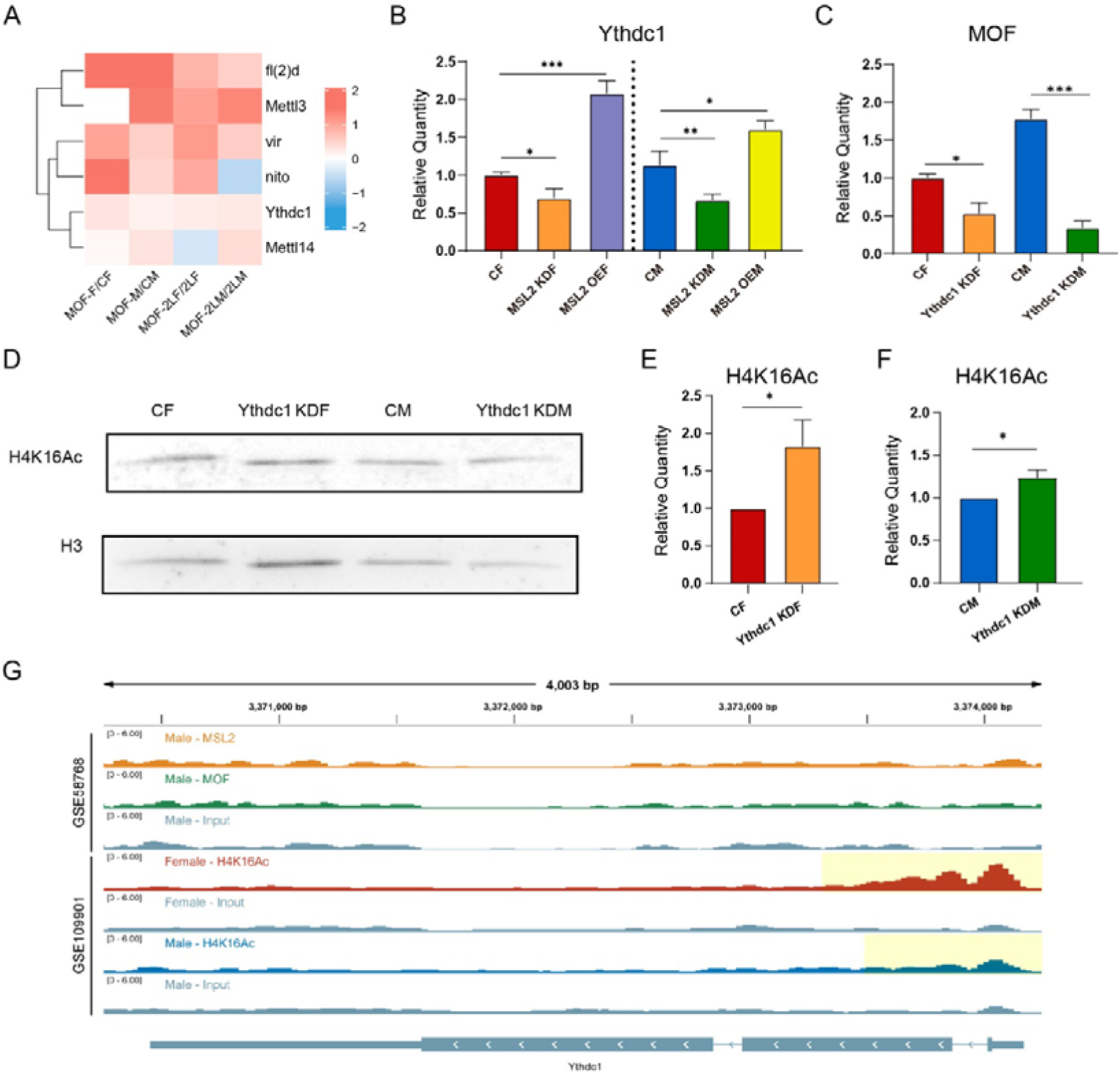
RNA m^6^A modification regulate histone acetyltransferase MOF and H4K16Ac. (A) Heatmap of the expression changes of m^6^A regulators in MOF overexpressing Drosophila larvae. The color of the heatmap represents log2(ratio). CF, wildtype female control; CM, wildtype male control; MOF-F, MOF-overexpressed female; MOF-M, MOF-overexpressed male; 2LF, trisomy 2L female; 2LM, trisomy 2L male; MOF-2LF, MOF-overexpressed trisomy 2L female; MOF-2LM, MOF-overexpressed trisomy 2L male. (B) RT-qPCR analysis of mRNA levels of Ythdc1 in the heads of MSL2 transgenic Drosophila. MOF KDF, MOF neural-knockdown female; MOF KDM, MOF neural-knockdown male; MOF OEF, MOF-overexpressed female; MOF OEM, MOF-overexpressed male (C) RT-qPCR analysis of mRNA levels of MOF in the heads of Ythdc1 knockdown Drosophila. Ythdc1 KDF, Ythdc1 neural-knockdown female; Ythdc1 KDM, Ythdc1 neural-knockdown male. (D) Western blot analysis of H4K16Ac in Drosophila. (E,F) Relative quantification of H4K16Ac in wild type and Ythdc1 knockdown adult Drosophila based on western blot (N = 5). Ythdc1 KDF, Ythdc1-knockdown female; Ythdc1 KDM, Ythdc1-knockdown male. (G) Genome browser example of Ythdc1 for indicated ChIP-seq data. Signals were displayed as CPM values. The gene architecture was shown at the bottom (only one representative transcript isoform was shown). The yellow shaded area indicates the presence of the H4K16Ac peaks..

Next, we investigated whether the m^6^A reader protein Ythdc1 would also have an effect on the levels of H4K16Ac in Drosophila. To this end, we employed western blot analysis to assess the expression patterns of H4K16Ac in Ythdc1 knockdown Drosophila adults (Figure 8D). The subsequent quantitative analysis revealed a significant upregulation of H4K16Ac in both female and male (Figure 8E,F). Furthermore, to substantiate our findings, we conducted polytene chromosome immunofluorescence in third instar larvae (Figure8—figure supplement 1G). Quantitative analysis of these assays in Ythdc1 knockdown Drosophila showed that the changes of H4K16Ac levels also showed sexual dimorphism (Figure 8—figure supplement 1H). In males, knockdown of Ythdc1 led to a decrease in H4K16Ac level; whereas in females, knockdown of Ythdc1 did not affect the level of H4K16Ac (Figure 8—figure supplement 1H). These observed changes in H4K16Ac were consistent with the pattern of changes in MOF protein in Ythdc1 knockdown Drosophila strains, suggesting that m^6^A modification may affect H4K16Ac levels through the mediation of MOF. Overall, while the alterations in H4K16Ac levels are not uniformly consistent between larvae and adult Drosophila, these findings nonetheless demonstrate that the knockdown of Ythdc1 has a significant impact on the expression levels of H4K16Ac at different stages of development and imply a potential relationship between H4K16Ac with m^6^A modification.

We analyzed two ChIP-seq datasets (GSE109901 and GSE58768) to study whether m^6^A regulator genes (especially Ythdc1) are targets of DCC components and H4K16Ac. According to the results, most of the m^6^A regulator genes, including Ythdc1, contain H4K16Ac peaks in both sexes, all of which are located in the 5’ regions (Figure 8G; Figure 8—figure supplement 2); except that Ime4 shows sexual dimorphism and only contains H4K16Ac peak in females. On the other hand, analysis of ChIP-seq data of MSL2 and MOF in male Drosophila showed that most of MSL2 and MOF peaks were located on the X chromosome (99.1% of MSL2 peaks and 61.6% of MOF peaks), which may be due to the fact that MSL2 and MOF are mostly tethered to the X chromosome by MSL complex under physiological conditions (Bashaw and Baker 1995; Kelley et al., 1995; Kind et al., 2008; Conrad et al., 2012). Therefore, there is no MSL2 and MOF peak near the m^6^A regulator genes which located on the autosomes (Figure 8H; Figure 8—figure supplement 2). These results showed that there is a direct relationship between m^6^A regulators and H4K16Ac, but there is no evidence that m^6^A regulator genes are direct targets of DCC components. MSL2 and MOF may thereby interact with m^6^A regulators in other ways.

To further study whether unbalanced genomes are involved in the interaction between m^6^A and histone acetylation modification, we also analyzed RNA sequencing data from trisomy 2L Drosophila strains overexpressing MOF. The data showed that overexpression of MOF in trisomy 2L resulted in significant changes in the expression of m^6^A regulators, with a trend different from that observed in diploids (Figure 8A). These results suggest that genomic imbalance might affect the interaction between MOF and m^6^A regulators to some extent, and the potential mechanisms require further investigation.

## Discussion

As an emerging epigenetic modification, RNA m^6^A methylation has been found to be involved in almost all aspects of RNA fate and metabolism (Gilbert et al., 2016; Yang et al., 2018). RNA m^6^A modification is also closely related to the development of organisms and a variety of human diseases (Barbieri et al., 2017; Zhao et al., 2017; Pinello et al., 2018; Ma et al., 2019; Liu et al., 2021; Shafik et al., 2021). However, the roles of m^6^A methylation in development and gene expression of aneuploidy have not been studied yet. Aneuploid variation is usually more detrimental than changes of the entire chromosome set due to genomic imbalance (Birchler and Veitia, 2007, 2012). The global changes of gene regulatory networks in unbalanced genomes involves various epigenetic mechanisms, including histone modification, chromatin remodeling, lncRNAs, microRNAs, etc. (Birchler, 2016; Zhang et al., 2021a, 2023; Shi et al., 2022). This study demonstrated that the expression of m^6^A components was altered under genomic imbalance, leading to dynamic changes in the entire methylome. Potential intermediaries by which m^6^A modification could affect trans regulation and achieve dosage compensation in aneuploid Drosophila were also investigated, such as dosage-sensitive modifiers, alternative splicing events, and the MSL complex.

Our experiments show that the expression levels of most m^6^A component genes are significantly down-regulated in aneuploid Drosophila larvae (Figure 1A; Figure 1—figure supplement 1C). Depletion of m^6^A components interferes with the development of animals and plants, especially leading to impaired self-renewal and differentiation of embryonic stem cells, defects in embryonic development, and even early embryonic lethality (Granadino et al., 1990; Horiuchi et al., 2006; Raffel et al., 2007; Luo et al., 2014; Wang Y et al., 2014; Geula et al., 2015). We demonstrated by TSA-FISH that appropriate temporal and spatial specific distributions of the transcripts for m^6^A components are vital during Drosophila embryogenesis (Figure 1C-F; Figure 1—figure supplement 2). The abnormal expression of m^6^A-related genes may affect the development of aneuploid embryos.

In previous studies, the abundance of m^6^A modification is usually positively correlated with the number of m^6^A peaks and m^6^A-marked genes (Luo et al., 2014; Zhu et al., 2023). However, our results obviously did not conform to this rule, with higher m^6^A abundance and fewer MeRIP-Seq peaks in aneuploids (Figure 2A-C). This reflects the complexity and heterogeneity of m^6^A modification. We suspect that in aneuploidy many RNAs may be lost that are methylated at a low level in wildtype, and possess a higher proportion of highly methylated RNAs. Analysis of the expression levels at each m^6^A site confirmed our hypothesis (Figure 2D). Thus, this phenomenon represents an imbalance in m^6^A methylation caused by aneuploidy. In addition, it is worth noting that due to the limitation of the larval samples, our detection of the overall abundance of m^6^A in aneuploidy is carried out for total RNA, including all types of RNA such as mRNA, lncRNA, and rRNA, and may be slightly different from detection of mRNA only. However, according to the results of m^6^A abundance detection in Drosophila adult heads, the enrichment or non-enrichment of mRNA from total RNA did not make a substantive difference in the results.

The distribution of m^6^A sites on gene features, m^6^A consensus motifs, biotypes of m^6^A-marked genes, enriched functions and pathways of methylated genes in aneuploidies are similar to those in wildtype. We also found that genes highly methylated in all genotypes are enriched for transcription factors (Figure 2I). It is consistent with previous studies suggesting that transcription regulatory genes may be preferentially targeted by m^6^A (Kan et al., 2017).

We also analyzed the characteristics of DMPs and DMP-associated genes in trisomies. By observing the distributions of MeRIP-Seq reads around DMP sites, it can be found that the densities of m^6^A-marked reads on chromosome 2L are relatively higher in trisomy 2L females and males (Figure 3F,G). Meanwhile, metafemales with triple X have a higher density of m^6^A-marked reads on chromosome X (Figure 3H). Previous studies have found that there is dosage compensation for cis-genes in autosomal and sex chromosome aneuploid Drosophila, and the expression of most genes on varied chromosomes approaches diploid levels (Sun et al., 2013b,c). Therefore, the up-regulation of m^6^A levels is not directly caused by the increased number of chromosomes. A recent study found that the selective enrichment of m^6^A methylation may play a role as a transcript degradation signal in dosage compensation in mammals (Rücklé et al., 2023).

Therefore, we speculate that changes in m^6^A levels in aneuploid Drosophila may affect its dosage compensation and inverse dosage effect by regulating the stability of the transcripts. In addition, more up-regulated DMPs are detected in trisomy 2L males and metafemales, while trisomy 2L females have more down-regulated DMPs (Figure 3A-D). Combined with the fact that the survival rate of trisomy 2L female larvae is relatively lower than that of the other two trisomies, it can be speculated that the regulation of DMPs may affect the survival and development of aneuploid Drosophila.

Gene expression in unbalanced genomes is extensively modulated (Sun et al., 2013b,c; Hou et al., 2018; Raznahan et al., 2018; Shi et al., 2021; Yang et al., 2021; Zhang et al., 2021b; San Roman et al., 2023). Previous studies have analyzed the transcriptome data of autosomal and sex chromosome trisomic Drosophila, and found that most of the genes on the triple chromosomes were compensated, and the ratios of their expression levels to wildtype were approximately 1; meanwhile, the expression of genes on other chromosomes was close to two-thirds of that of the wildtype, which was called an inverse dosage effect (Sun et al., 2013b,c; Zhang et al., 2021b). We investigated the relationships between genes with different m^6^A methylation status and genes modulated by classical dosage-related effects (Figure 5F-H). The results showed that for aneuploid females, genes methylated in both trisomy and control are significantly enriched in dosage compensated cis genes and inverse dosage effect trans genes, and genes not methylated at all are enriched in dosage effect genes. However, in aneuploid males, methylated genes are associated with gene dosage effect (Figure 5F-H). We also found that the proportion of 5’UTR m^6^A peaks on dosage compensation and inverse dosage effect genes are increased in aneuploid females (Figure 5I-K). These results provide evidence of sexual dimorphism in the relationships between RNA m^6^A modification and dosage-related effects. m^6^A located in the 5’UTR generally shows higher tissue specificity and richer dynamic changes, and may have unique regulatory functions (Dominissini et al., 2012; Meyer and Jaffrey, 2014; Gilbert et al., 2016). The increased proportion of m^6^A in the 5’UTR of genes with classical dosage-related effects in aneuploidies suggests that m^6^A modification may be involved in the regulation of dosage-dependent genes.

Dosage-related effects of aneuploidy are thought to be caused by dosage-sensitive genes (Shi et al., 2021; Yang et al., 2021). Some dosage-sensitive regulators have been identified in Drosophila, and changes in the dosage of individual regulatory genes can mimic the effects of aneuploidy in the whole genome (Birchler et al., 2001; Xie and Birchler, 2012; Zhang et al., 2021b). Most dosage-sensitive regulators are transcription factors, signal transduction components, and chromatin proteins, which have in common being members of macromolecular complexes or having multicomponent interactions (Birchler et al., 2001; Birchler and Veitia, 2007). We studied the m^6^A modification of dosage-dependent genes and their protein-protein interaction networks in aneuploid Drosophila (Figure 5—figure supplement 1). The results showed that there are complex interactions between dosage-sensitive regulators and differentially expressed genes. Among them, most of the DEGs located on the unvaried chromosomes are down-regulated, indicating that the dosage-sensitive regulators mainly have negative effects on trans target genes. There are more regulatory genes with up-regulated DMPs than down-regulated DMPs in all three aneuploidies, and many of the regulators with up-regulated methylation are connected with interactors that have down-regulated expression.

We comprehensively analyzed the relationships among RNA m^6^A modification, gene expression levels, and alternative splicing. The data showed that differential m^6^A methylation under genomic imbalance appeared to be more closely associated with differential alternative splicing (Figure 5A-C; Figure 6A-C). This phenomenon is reasonable because the mutation or knockout of Ime4, dMettl14, fl(2)d, and Ythdc1 has been found to affect a large number of alternative splicing events in Drosophila, and fl(2)d itself is thought to encode a splicing factor (Penn et al., 2008; Haussmann et al., 2016; Lence et al., 2016). A small set of transcripts are simultaneously differentially m^6^A methylated, differentially expressed, and differentially alternative spliced in aneuploidies, including the transcription factor su(Hw), which may be a dosage-sensitive regulator (Figure 6G-H). Besides, m^6^A modification in the 5’UTR regions may play a special role in some differential alternative splicing events (Figure 6—figure supplement 1F-I).

RNA m^6^A modification in Drosophila has been shown to be involved in the alternative splicing of Sxl (Lence et al., 2017). The deletion of some m^6^A components in females will result in a reduction of female-specific splicing of Sxl and an increase of the inclusion of the third exon, along with phenotypic sexual transformation (Haussmann et al., 2016; Lence et al., 2016; Kan et al., 2017). We found that Sxl transcripts of aneuploid Drosophila are both differentially m^6^A methylated and differentially spliced (Figure 7A-B). Among multiple alternative splicing events, trisomy 2L females have a higher level of the third exon inclusion compared with wildtype females (Figure 7B), which may be related to the differential methylation at 5’UTR of the Sxl transcripts.

Some studies proposed that the deletion of RNA m^6^A methyltransferase Ime4 harms the survival of female Drosophila because of the insufficient inhibition of msl-2 caused by decreased Sxl levels, which in turn leads to up-regulation of X-linked genes (Haussmann et al., 2016). However, other studies have found that the expression of genes on the X chromosome in females with ectopic expression of MSL2 does not increase twofold, that is, the MSL complex does not directly mediate dosage compensation, and the global inverse dosage effect caused by the imbalance of sex chromosomes is the basis for dosage compensation (Birchler, 1981, 2016; Sun et al., 2013a). Therefore, the lethality of the lack of m^6^A in female Drosophila may not be directly caused by up-regulation of X-linked genes through ectopic assembly of MSL complex, and the specific reasons need to be further studied.

The functions of MSL complex in dynamic regulation of global gene expression in aneuploid genomes has been described (Zhang et al., 2021a). Here, we found that MSL2, a structure component of MSL complex, not only affects the expression of m^6^A regulators, but also influences the overall abundance of mRNA m^6^A in Drosophila (Figure 7C-F), proving a close relationship between the MSL complex and RNA m^6^A modification. In addition, under the condition of genomic imbalance, the relative expression of m^6^A components was also changed in larvae and embryos of MSL2-overexpressed trisomies (Figure 7G,H). Another component of the MSL complex, MOF, which mediates histone acetylation and transcriptional activation (Kind et al., 2008; Conrad et al., 2012), is also closely associated with the m^6^A reader Ythdc1. Overexpression of MOF increased the expression of Ythdc1 (Figure 8A,B), and in turn, knockdown of Ythdc1 influenced the expression of MOF and the level of H4K16Ac catalyzed by it (Figure 8C-F; Figure 8—figure supplement 1G,H). It is worth noting that the rationale behind the variable expression levels of H4K16Ac in Ythdc1 knockdown Drosophila across different developmental stages merits further investigation. These results demonstrate complicated interactions between RNA m^6^A methylation and the MSL complex in unbalanced genomes, which may affect the gene expression, sexual dimorphism, development and survival of aneuploid Drosophila.

## Materials and methods

### Key resources table

Softwares used in this study:

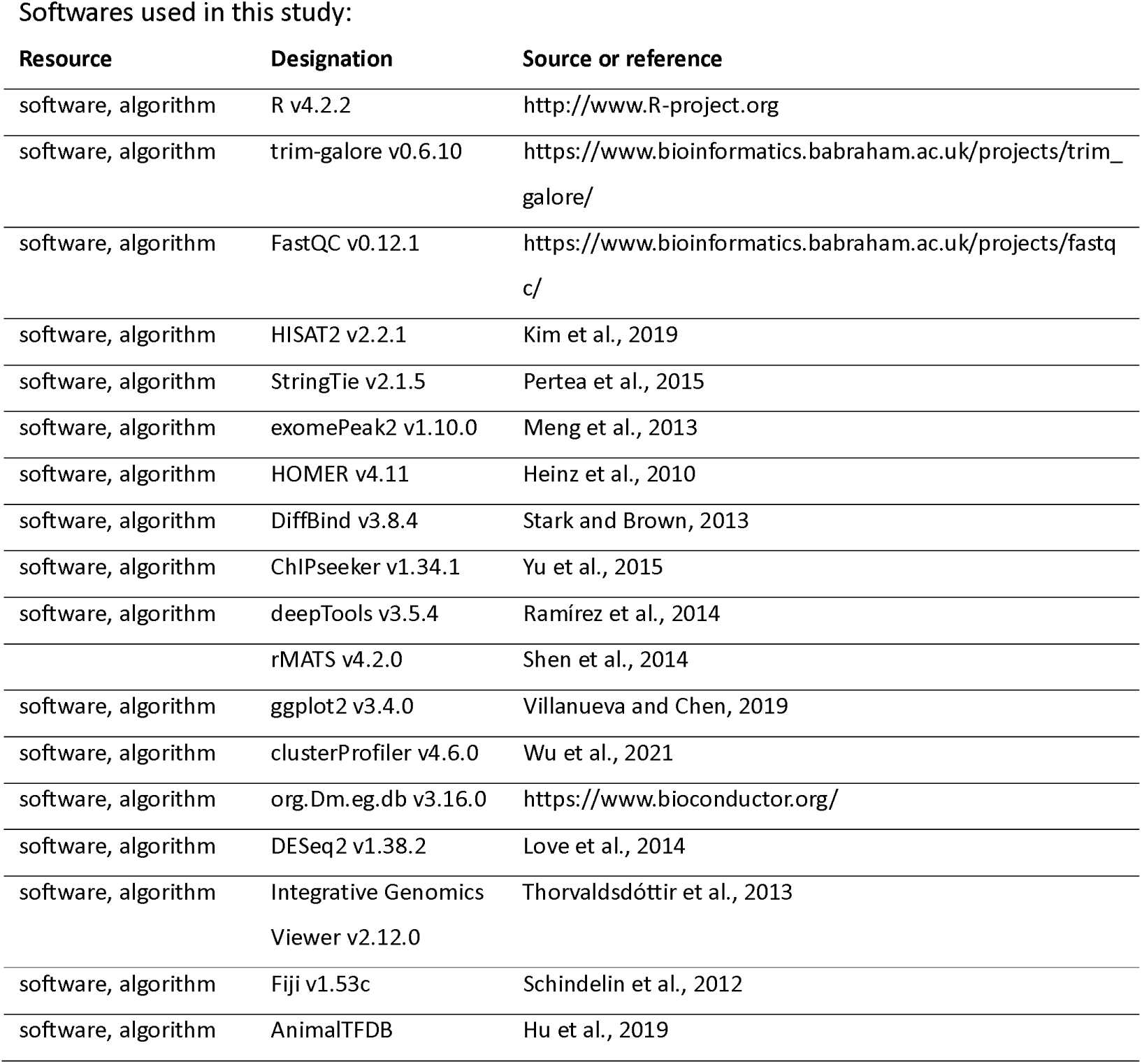
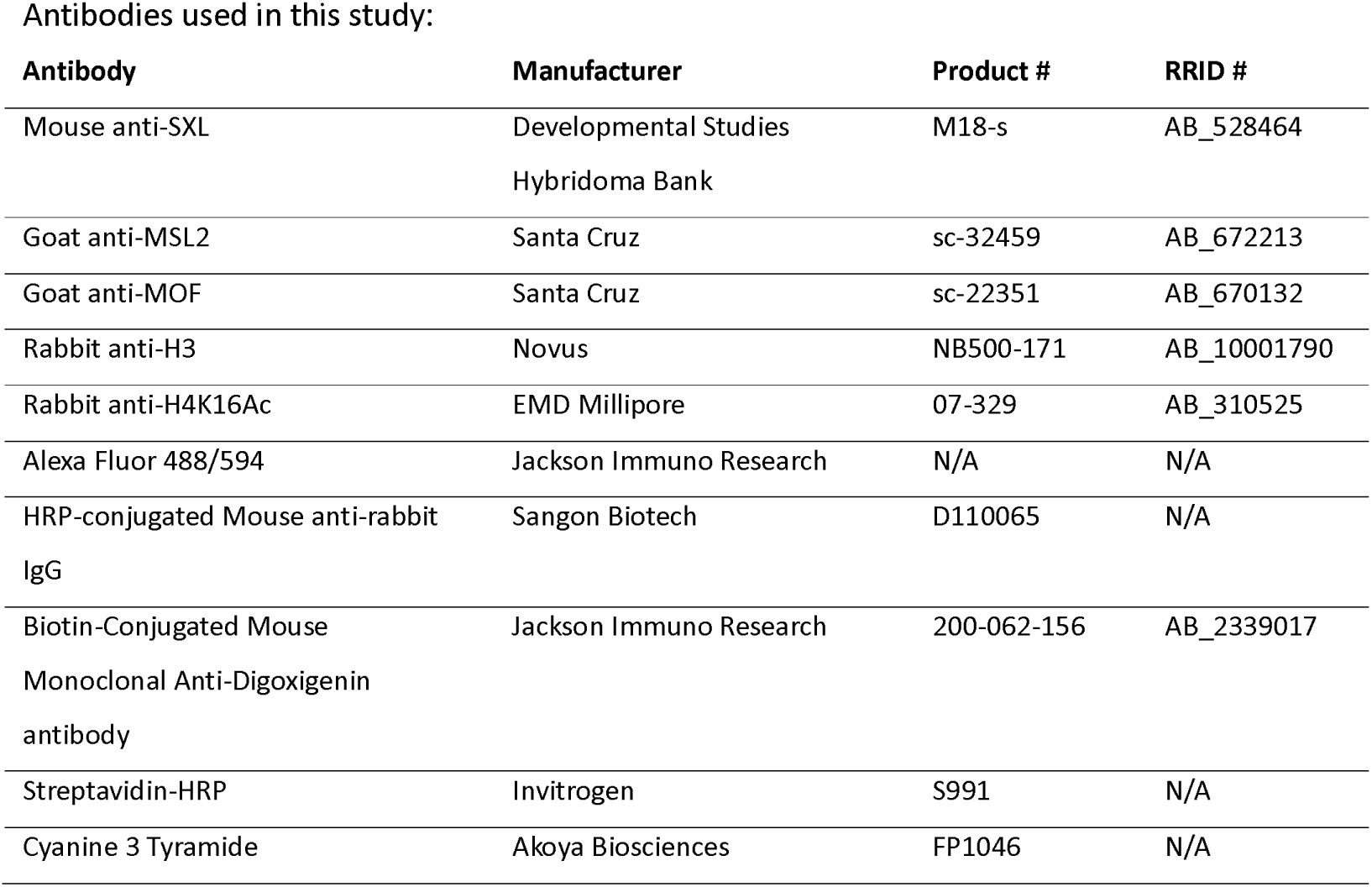

| Resource | Designation | Source or reference |
| --- | --- | --- |
| software, algorithm | R v4.2.2 | http://www.R-project.org |
| software, algorithm | trim-galore v0.6.10 | https://www.bioinformatics.babraham.ac.uk/projects/trim_galore/ |
| software, algorithm | FastQC v0.12.1 | https://www.bioinformatics.babraham.ac.uk/projects/fastqc/ |
| software, algorithm | HISAT2 v2.2.1 | Kim et al., 2019 |
| software, algorithm | StringTie v2.1.5 | Pertea et al., 2015 |
| software, algorithm | exomePeak2 v1.10.0 | Meng et al., 2013 |
| software, algorithm | HOMER v4.11 | Heinz et al., 2010 |
| software, algorithm | DiffBind v3.8.4 | Stark and Brown, 2013 |
| software, algorithm | ChIPseeker v1.34.1 | Yu et al., 2015 |
| software, algorithm | deepTools v3.5.4 | Ramírez et al., 2014 |
|  | rMATS v4.2.0 | Shen et al., 2014 |
| software, algorithm | ggplot2 v3.4.0 | Villanueva and Chen, 2019 |
| software, algorithm | clusterProfiler v4.6.0 | Wu et al., 2021 |
| software, algorithm | org.Dm.eg.db v3.16.0 | https://www.bioconductor.org/ |
| software, algorithm | DESeq2 v1.38.2 | Love et al., 2014 |
| software, algorithm | Integrative Genomics Viewer v2.12.0 | Thorvaldsdóttir et al., 2013 |
| software, algorithm | Fiji v1.53c | Schindelin et al., 2012 |
| software, algorithm | AnimalTFDB | Hu et al., 2019 |

Antibodies used in this study:
| Antibody | Manufacturer | Product # | RRID # |
| --- | --- | --- | --- |
| Mouse anti-SXL | Developmental Studies<br>Hybridoma Bank | M18-s | AB_528464 |
| Goat anti-MSL2 | Santa Cruz | sc-32459 | AB_672213 |
| Goat anti-MOF | Santa Cruz | sc-22351 | AB_670132 |
| Rabbit anti-H3 | Novus | NB500-171 | AB_10001790 |
| Rabbit anti-H4K16Ac | EMD Millipore | 07-329 | AB_310525 |
| Alexa Fluor 488/594 | Jackson Immuno Research | N/A | N/A |
| HRP-conjugated Mouse anti-rabbit<br>IgG | Sangon Biotech | D110065 | N/A |
| Biotin-Conjugated Mouse<br>Monoclonal Anti-Digoxigenin<br>antibody | Jackson Immuno Research | 200-062-156 | AB_2339017 |
| Streptavidin-HRP | Invitrogen | S991 | N/A |
| Cyanine 3 Tyramide | Akoya Biosciences | FP1046 | N/A |

### Drosophila stocks and genetic crosses

The Drosophila strains mentioned in this study were all maintained and crossed in our laboratory. The crossing methods have been described previously (Zhang et al., 2021a). Trisomy chromosome 2 left arm (2L) female and male third instar larvae were obtained from the cross of y; C(2L)dp; F(2R) bw females and Canton S males. The metafemale larvae were obtained from the cross of C(1)DX, ywf/winscy females and Canton S males. Aneuploidies overexpressing MSL2 were generated from the crosses of Drosophila with compound chromosomes and MSL2 homozygotes. The MSL2 transgene strain was constructed and validated in a previous study (Sun et al., 2013a). All Drosophila strains were cultured on cornmeal dextrose medium at 25°C. Genes and chromosomal balancers are described in Flybase (https://flybase.org/).

### RNA m A methylation quantification

Total RNA was extracted from Drosophila larvae using TRIzol Reagent (Invitrogen) and mRNA was isolated using the Dynabeads mRNA purification kit (Invitrogen, 61006) to detect the abundance of m^6^A. The relative quantification of m^6^A methylation was performed using EpiQuik m^6^A RNA Methylation Quantification Kit (Colorimetric) (Epigentek, NY, USA, Cat # P-9005). Specifically, 80 µl of Binding Solution was first added to each well of the plate, and 200 ng of total RNA or mRNA samples, 2 µl of Negative Control, or 2 µl of diluted Positive Control were added to the designed wells. Subsequently, the plate was incubated at 37°C for 90 min. After washing with Wash Buffer, 50 µl of Capture Antibody, 50 µl of Detection Antibody, and 50 µl of Enhancer Solution were added to each well in order, and wells were emptied before adding a new solution each time. Finally, 100 µl of Developer Solution and Stop Solution were added to each well away from light, and the absorbance was read on a microplate reader at a wavelength of 450 nm.

### m A methylated RNA immunoprecipitation sequencing (MeRIP-Seq)

The MeRIP-Seq service was provided by Cloudseq Biotech Inc. (Shanghai, China), and this technology was developed on the basis of published experimental methods (Meyer et al., 2012). In brief, Drosophila larvae of five genotypes, each with two biological replicates, were collected for sequencing. Ribosomal RNAs were removed from total RNA using Ribo-Zero rRNA Removal Kits (Illumina, USA). Immunoprecipitation of m^6^A RNA was performed using GenSeq® m^6^A RNA IP kit (GenSeq, Shanghai, China). The NEBNext® Ultra II Directional RNA Library Prep kit (New England Biolabs, USA) was used for RNA sequencing library construction. High-throughput sequencing was performed using Illumina NovaSeq 6000 sequencers with the paired-end 150 bp protocol.

### Analysis of MeRIP-Seq data

The raw sequencing data was first filtered by Trim Galore (version 0.6.10) (https://www.bioinformatics.babraham.ac.uk/projects/trim_galore/) to remove adapters and low-quality reads. The quality of the data was then assessed by FastQC (version 0.12.1) (https://www.bioinformatics.babraham.ac.uk/projects/fastqc/). Subsequently, clean reads were aligned to the Drosophila reference genome (Drosophila_melanogaster.BDGP6.32.dna.toplevel.fa, downloaded from the Ensembl database) using HISAT2 (version 2.2.1) (Kim et al., 2019). Next, The R package exomePeak2 (version 1.10.0) (Meng et al., 2013) was used for m^6^A peak calling (the screening criteria were log2FC ≥ 1, RPM.IP ≥ 0.5, and score ≥ 5). Motif analysis was performed using HOMER (version 4.11) (Heinz et al., 2010). Differentially methylated peaks were analyzed by the R package DiffBind (version 3.8.4) (Stark and Brown, 2013), which employs the DESeq2 (Love et al., 2014) method. m^6^A peaks with a p-value of 0.1 or less were considered as differentially methylated peaks (DMPs). ChIPseeker (version 1.34.1) (Yu et al., 2015) was used to annotate the peaks, and plotted some of the figures. The profiles and heatmaps of MeRIP-Seq reads around DMPs were generated by deepTools (version 3.5.4) (Ramírez et al., 2014). The signal distribution on the genome was visualized using the Integrative Genomics Viewer (IGV) software (version 2.12.0) (Thorvaldsdóttir et al., 2013).

### RNA sequencing data and analysis

The RNA-seq data of aneuploid Drosophila used in this article were generated in a previous study (Zhang et al., 2023) and can be downloaded from the Gene Expression Omnibus (GEO) database (GSE233534). Genome mapping of sequencing data and gene expression quantification were carried out through HISAT2-StringTie pipeline (Pertea et al., 2015; Kim et al., 2019). Differential alternative splicing events were identified by rMATS (version 4.2.0) program (Shen et al., 2014) (with the parameters of -t paired --readLength 150 --cstat 0.0001 --libType fr-firststrand). The method of generating the ratio distributions of gene expression changes was as described previously (Zhang et al., 2022). The plots were generated using ggplot2 (version 3.4.0) (Villanueva and Chen, 2019) in the R program (version 4.2.2). Differential expression analysis was performed by DESeq2 (version 1.38.2) (Love et al., 2014), and the threshold was set to adjusted p-value ≤ 0.05. The functional enrichment analysis was performed by ClusterProfiler (version 4.6.0) (Wu et al., 2021) based on org.Dm.eg.db (version 3.16.0) from Bioconductor (https://www.bioconductor.org/). The Kyoto Encyclopedia of Genes and Genomes (KEGG) pathway data was obtained from the network (https://www.kegg.jp/). The list of transcription factors of Drosophila was downloaded from AnimalTFDB (Hu et al., 2019). Protein-protein interaction (PPI) relationships were obtained from the STRING database (Szklarczyk et al., 2019).

### Real-time quantitative PCR

Total RNA of Drosophila whole larvae, larval brains, or adult heads was extracted using TRIzol Reagent (Invitrogen), and reverse transcription was done with TransScript one-step gDNA Removal and cDNA Synthesis SuperMix (TransGen Biotech). The sequences of primers designed for RT-qPCR were listed in Figure 1—figure supplement 1A. The usability of these primers was verified by agarose gel electrophoresis (Figure 1—figure supplement 1B). The real-time PCR was performed with TransStart Tip Green qPCR SuperMix (+ Dye II) (TransGen Biotech) using ABI QuantStudio 6 Flex Real-Time PCR System. Relative quantification of gene expression was determined using the 2^-ΔΔCt^ method.

### Fluorescence in situ hybridization (FISH) of Drosophila embryos

Collection of Drosophila embryos and fluorescence in situ hybridization were performed as previously described (Jandura et al., 2017; Zhang et al., 2021a). The probe primers containing flanking T7 promoter elements were listed in Figure 1—figure supplement 1D. After PCR amplification and in vitro transcription, their products were examined by agarose gel electrophoresis (Figure 1—figure supplement 1E). Digoxygenin (DIG)-labelled antisense RNA probes were hybridized to transcripts of interest, and then detected using a succession of anti-DIG antibody conjugated to biotin, streptavidin conjugated to horseradish peroxidase (HRP) and fluorescently conjugated tyramide. This hierarchical process can greatly enhance the probe signals and is referred to as a tyramide signal amplification (TSA) system. All images were acquired with a Zeiss LSM880 laser confocal fluorescence microscope using ZEN software. The same probes for relative fluorescence intensity analysis were photographed using the same parameters. Fluorescence images were processed and analyzed using Fiji (version 1.53c) (Schindelin et al., 2012).

### Polytene chromosomes immunostaining

Salivary gland chromosomes immunostaining was performed as previously described (Zhang et al., 2021a). Briefly speaking, the salivary glands from third-instar larvae were first fixed in 3.7% formaldehyde for 1 min, and then dissociated with 50% acetic acid for 5 min. Polytene chromosomes were treated with the following primary antibodies at a dilution of 1:100: anti-SXL (Developmental Studies Hybridoma Bank, M18-s), anti-MSL2 (Santa Cruz, sc-32459), anti-MOF (Santa Cruz, sc-22351), and anti-H4K16Ac (EMD Millipore, 07-329). Finally, fluorescence-conjugated secondary antibodies (Alexa Fluor 488 and Alexa Fluor 594, Jackson Immuno Research) were used for detection.

### Histone extraction and Western Blotting

Drosophila adults were ground and lysed in Triton Extraction Buffer (PBS at pH 7.4 with 0.5% Triton X-100, 0.5 mM phenymethylsulfonyl fluoride, and 0.02% sodium butyrate) and histones were acid-extracted in 0.2 N HCl overnight. Acid-extracted histones were then run on a 12% Tris-Glycine gel and blotted onto a PVDF membrane. Antibodies for Histone H3 (NB500-171, Novus) and H4K16Ac (07-329, Sigma) were all incubated at a dilution of 1:1000 in 5% skim milk powder solution. Westerns blots were imaged and protein levels quantified using the ImageJ software.

## Acknowledgements

This work was supported by National Natural Science Foundation of China (Grant No. 32070566) to L.S. We thank James A.Birchler for revising the manuscript.

## Additional information

## Funding

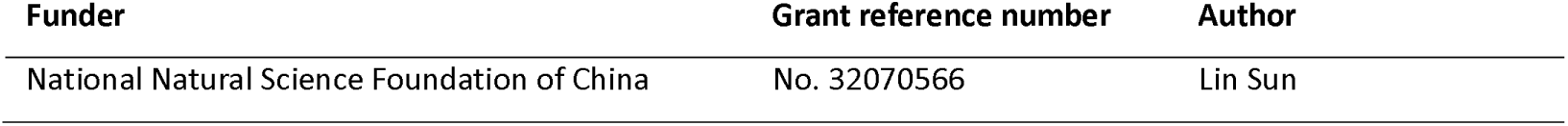

## Author contributions

Shuai Zhang: Methodology, Investigation, Data Curation, Formal analysis, Writing - Original Draft, Writing - Review & Editing, Visualization. Ruixue Wang, Methodology, Investigation, Validation, Writing - Original Draft, Writing - Review & Editing, Visualization. Kun Luo, Methodology, Investigation, Validation, Writing - Review & Editing, Visualization. Shipeng Gu, Methodology, Investigation, Writing - Review & Editing, Visualization. Xinyu Liu, Methodology, Investigation, Validation, Writing - Original Draft. Junhan Wang: Methodology, Investigation, Formal analysis. Ludan Zhang: Methodology, Investigation, Validation. Lin Sun: Conceptualization, Writing - Original Draft, Writing - Review & Editing, Supervision, Project administration, Funding acquisition. All authors have read and approved the final manuscript.

## Additional files Supplementary files

Supplementary File 1

## Data availability

All raw and processed sequencing data generated in this study have been submitted to the NCBI Gene Expression Omnibus (GEO; https://www.ncbi.nlm.nih.gov/geo/) under accession number GSE253401. The previously published RNA-seq data used in this study are available at the GEO repository under the accession number GSE233534. The previously published ChIP-seq data used in this study are available at the GEO repository under the accession number GSE109901 and GSE58768.

The following datasets were generated:

| Author(s) | Year | Dataset title | Dataset URL | Database and Identifier |
| --- | --- | --- | --- | --- |
| Zhang S, Liu X, Wang R, Wang J, Zhang L, Sun L | 2024 | Dynamics and regulatory roles of RNA m<sup>6</sup>A methylation in unbalanced genomes | https://www.ncbi.nlm.nih.gov/geo/query/acc.cgi?acc=GSE253401 | NCBI Gene Expression Omnibus, GSE253401 |

The following previously published datasets were used:

| Author(s) | Year | Dataset title | Dataset URL | Database and Identifier |
| --- | --- | --- | --- | --- |
| Zhang S, Wang R, Zhu X, Zhang L, Liu X, Sun L | 2024 | Characteristics and expression of lncRNA in *Drosophila* aneuploidy | https://www.ncbi.nlm.nih.gov/geo/query/acc.cgi?acc=GSE233534 | NCBI Gene Expression Omnibus, GSE233534 |
| Renschler G,<br>Richard G,<br>Valsecchi CIK,<br>Toscano S et al. | 2018 | Facultative dosage<br>compensation of<br>developmental genes on<br>autosomes in *Drosophila*<br>and mammals | https://www.ncbi.nlm.nih.gov/geo/query/acc.cgi?acc=GSE109901 | NCBI Gene Expression<br>Omnibus, GSE109901 |
| Figueiredo ML,<br>Kim M, Philip P,<br>Allgardsson A,<br>Stenberg P,<br>Larsson J | 2014 | Non-coding *roX* RNAs<br>prevent the binding of<br>the MSL-complex to<br>heterochromatic regions | https://www.ncbi.nlm.nih.gov/geo/query/acc.cgi?acc=GSE58768 | NCBI Gene Expression<br>Omnibus, GSE58768 |

## Figure supplements

**Figure 1—figure supplement 1.**
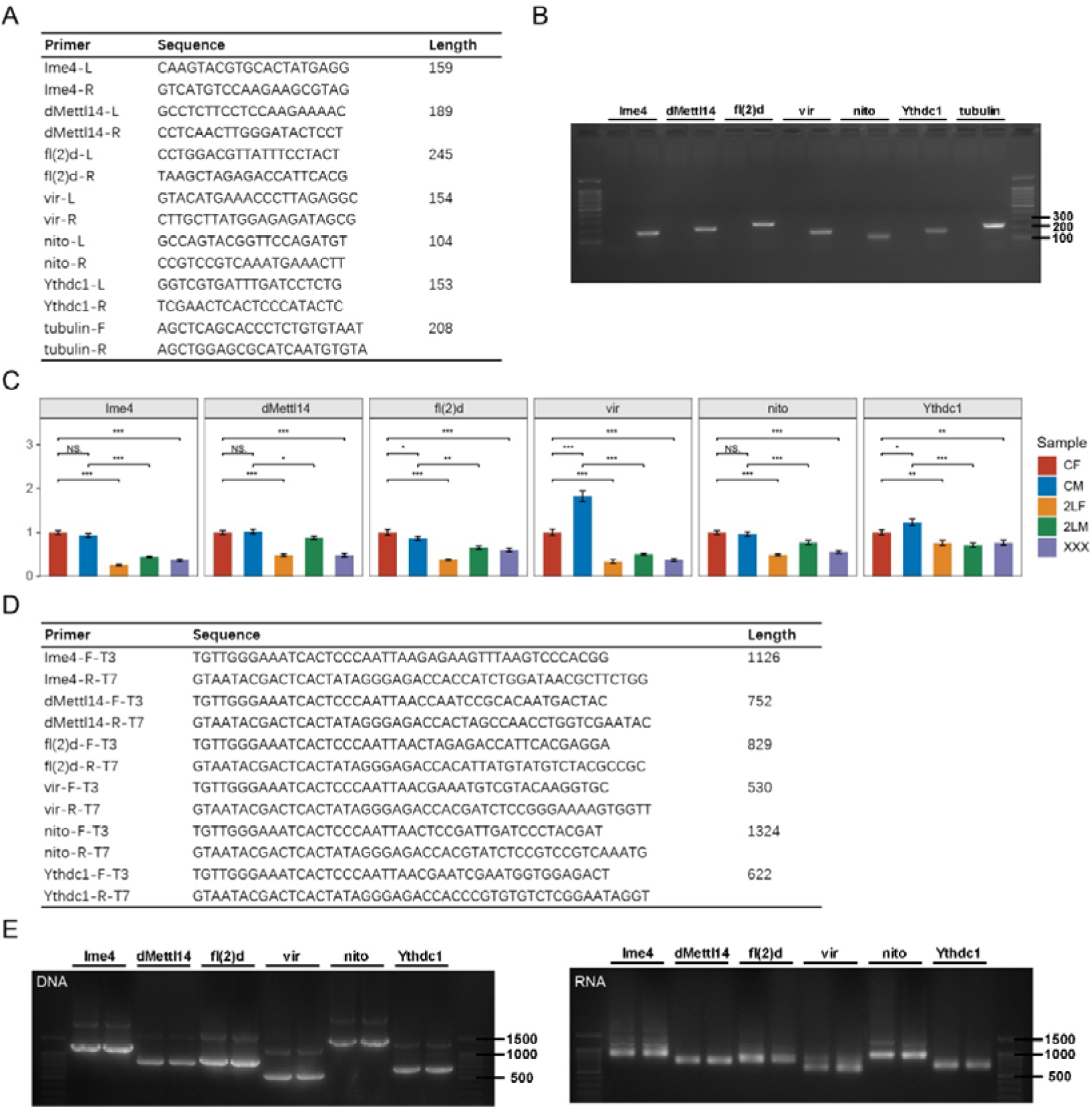
Primers of m^6^A methyltransferases and reader protein for RT-qPCR and TSA-FISH. (A) The sequences of RT-qPCR primers. (B) Agarose gel electrophoresis validation of qPCR primers. The left lane of each gene is negative control with primers only and no DNA substrate. (C) RT-qPCR analysis of mRNA levels of m^6^A methyltransferases and reader protein in the brains of wildtype and trisomy Drosophila larvae. CF, wildtype female control; CM, wildtype male control; 2LF, trisomy 2L female; 2LM, trisomy 2L male; XXX, metafemale. Student’s t test *p < 0.05, **p < 0.01, ***p < 0.001. (D) The sequences of FISH primers. (E) The sizes of the products of the FISH primers after PCR amplification and in vitro transcription were tested by agarose gel electrophoresis. 2L, chromosome 2 left arm; RT-qPCR, Real-time quantitative PCR; TSA-FISH, Tyramide signal amplification-based fluorescence in situ hybridization.

**Figure 1—figure supplement 2.**
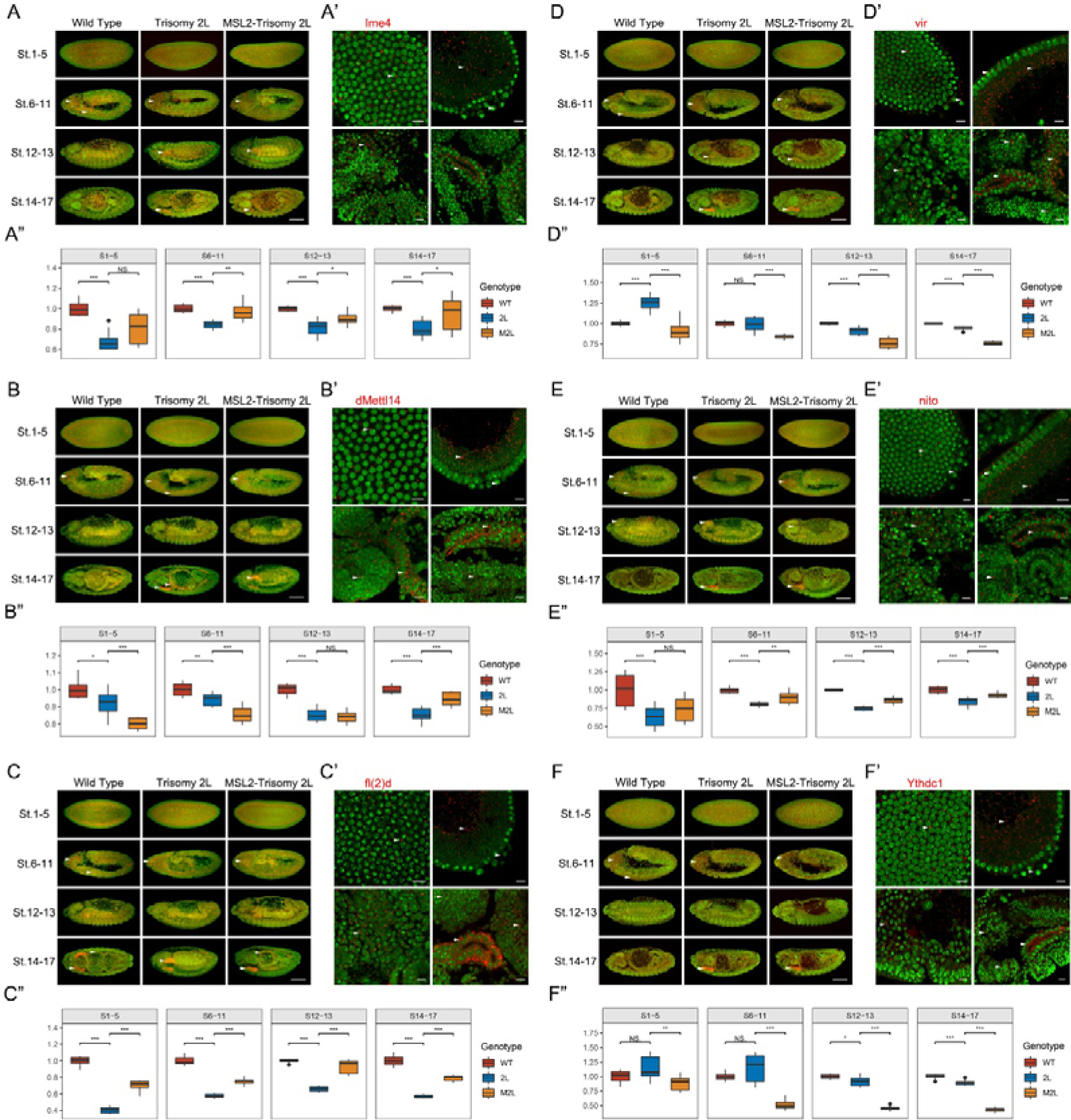
Embryo TSA-FISH of m^6^A components in wildtype, trisomy, and MSL2-overexpressed trisomy Drosophila. The images represent m^6^A component Ime4 (A-A”), dMettl14 (B-B”), fl(2)d (C-C”), vir (D-D”), nito (E-E”), and Ythdc1 (F-F”), respectively. (A-F) Gene expression patterns in entire embryos. The genotypes of the samples were shown above, and the development stages (St.) were shown in the left of the pictures. The red pseudocolor represents signal from the probes, and green is the signal of nucleus. Arrowheads indicate regions where the probe signal are enriched. Scale bars, 100 μm. (A’-F’) RNA subcellular location patterns of m^6^A components. Arrowheads indicate probe signals. Scale bars, 10 μm. (A”-F”) The expression levels of m^6^A component genes represented as relative fluorescence intensity of the probes compared with DAPI. The expression of wildtype embryos was set as base line. WT, wildtype; 2L, trisomy 2L; M2L, MSL2-overexpressed trisomy 2L. Sample size = 10. Student’s t test *p < 0.05, **p < 0.01, ***p < 0.001.

**Figure 2—figure supplement 1.**
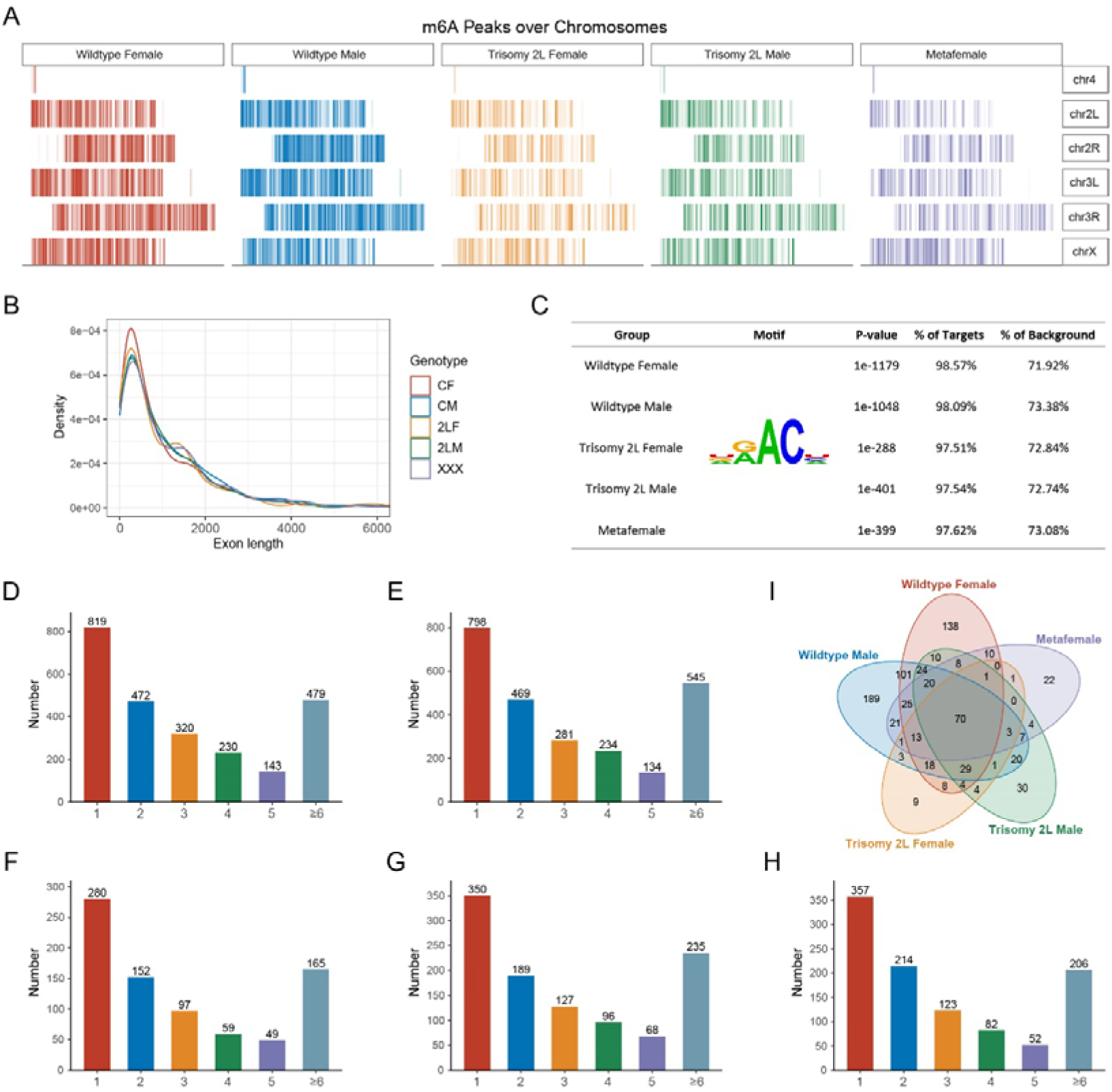
Characteristics of m^6^A-modified genes in wildtype and aneuploid Drosophila. (A) Distribution of m^6^A peaks along each chromosome. (B) Kernel density plot showing the distribution of exon lengths of m^6^A-modified genes. CF, wildtype female control; CM, wildtype male control; 2LF, trisomy 2L female; 2LM, trisomy 2L male; XXX, metafemale. (C) Enrichment analysis performed on the known m^6^A consensus motif DRACH (D = G/A/U, R = G/A, H = U/A/C). (D-H) Statistics of the number of m^6^A peaks on m^6^A-modified genes in wildtype females (D), wildtype males (E), trisomy 2L females (F), trisomy 2L males (G), and metafemale (H). (I) Venn diagram showing the number of m^6^A-modified genes with more than five m^6^A peaks.

**Figure 2—figure supplement 2.**
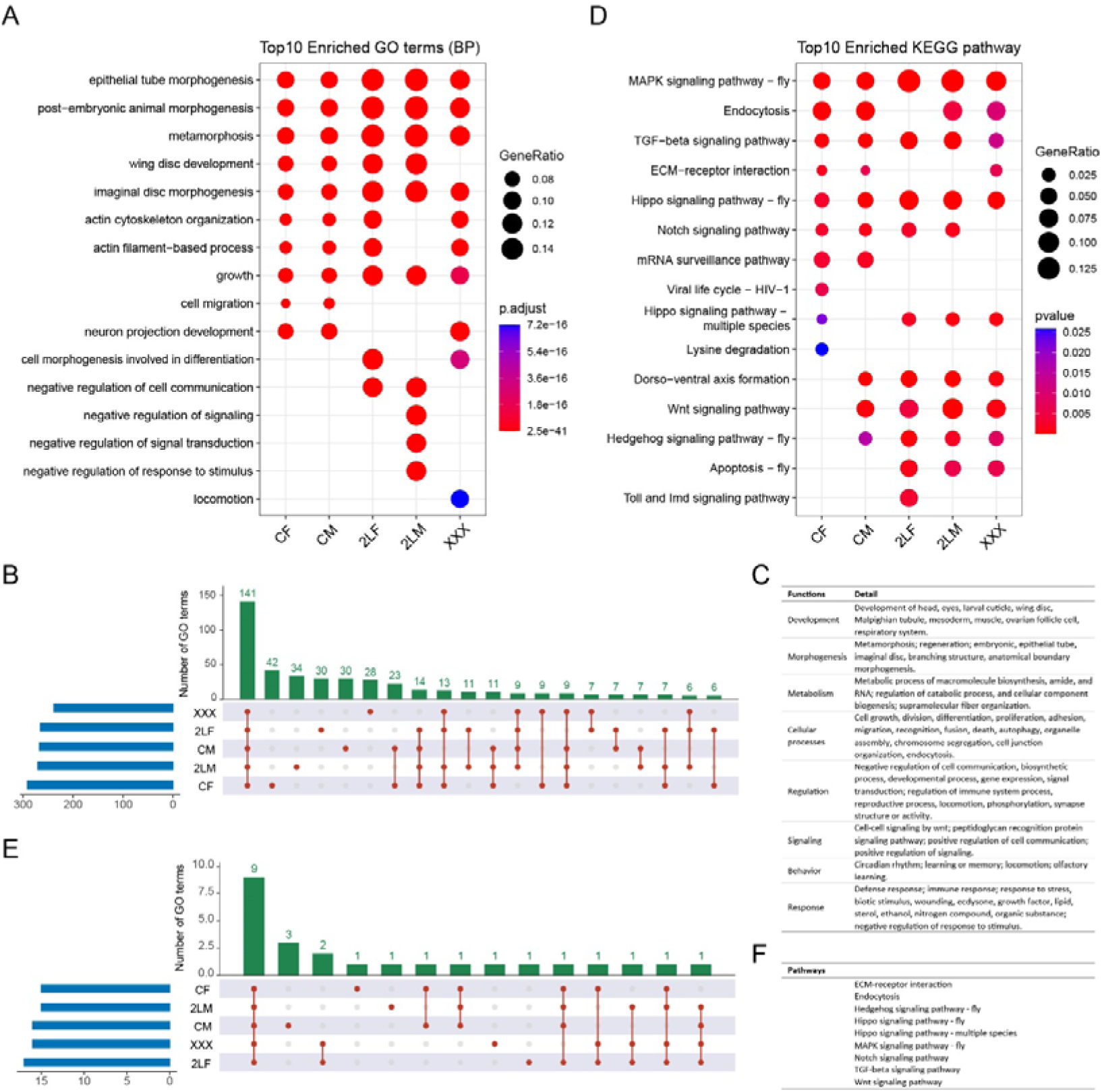
Functional and pathway analysis of m^6^A-modified genes in wildtype and aneuploid Drosophila. (A) Functional enrichment analysis of m^6^A-modified genes. Top 10 enriched GO terms (Biological Process) with adjusted p-value < 0.05 in each sample were shown. (B) UpSet plot showing the shared sets of enriched GO terms of m^6^A-modified genes in different groups. (C) Summary of the common functions enriched by m^6^A-modified genes in all genotypes. (D) KEGG pathway enrichment analysis of m^6^A-modified genes. Top 10 enriched pathways with p-value < 0.05 in each sample were shown. (E) UpSet plot showing the shared sets of enriched KEGG pathways of m^6^A-modified genes in different groups. (F) The pathways enriched in all genotypes were listed. CF, wildtype female control; CM, wildtype male control; 2LF, trisomy 2L female; 2LM, trisomy 2L male; XXX, metafemale; GO, Gene Ontology; BP, Biological Process; KEGG, Kyoto Encyclopedia of Genes and Genomes.

**Figure 3—figure supplement 1.**
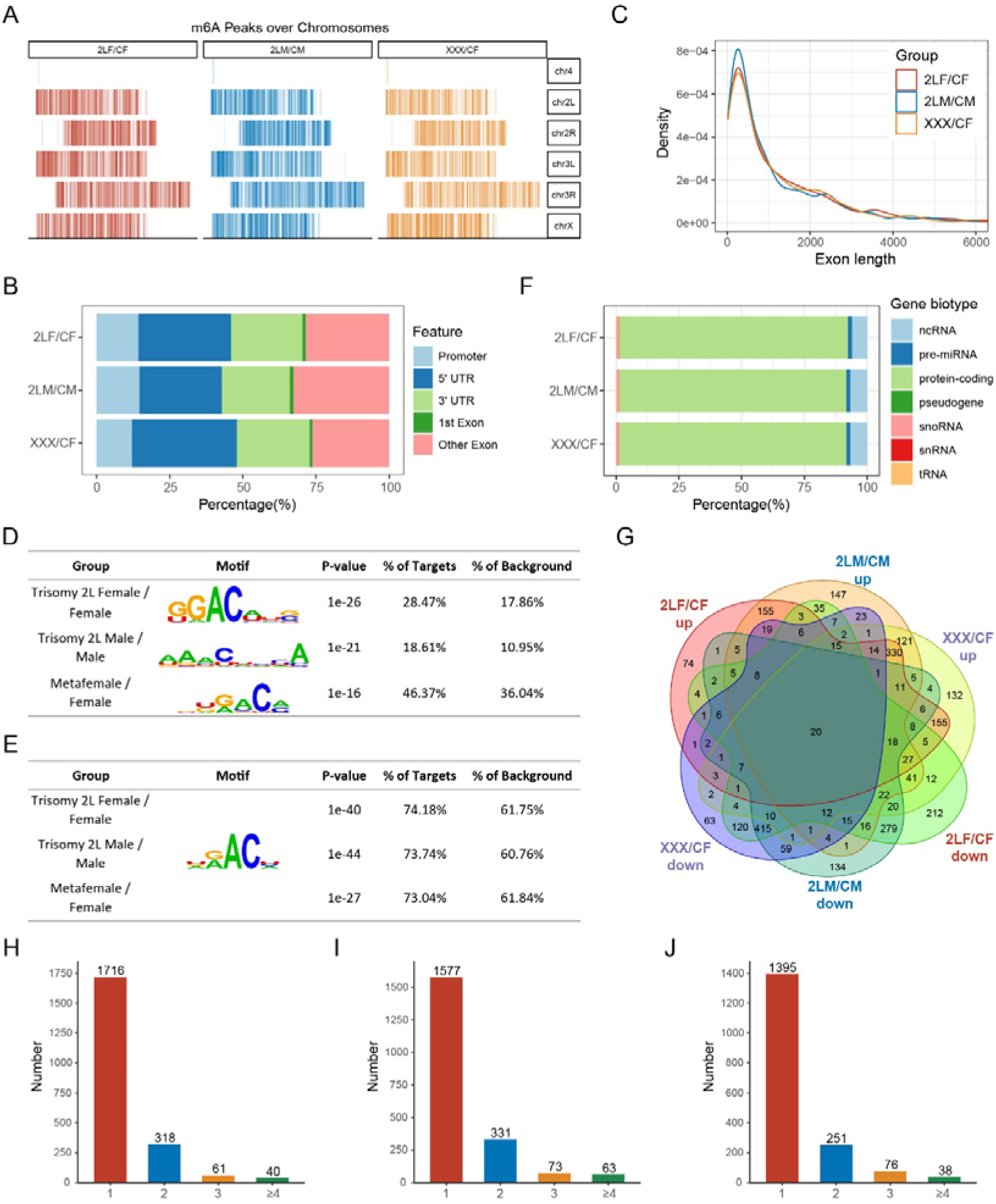
Characteristics of differentially methylated peak (DMP) associated genes. (A) Distribution of differentially methylated peaks (DMPs) along each chromosome. (B) Percentages of DMPs localized on different gene features. (C) Kernel density plot showing the distribution of exon lengths of DMP-associated genes. (D) The top enriched motifs with de novo motif analysis of DMPs in each comparison. (E) Enrichment analysis performed on the known m^6^A consensus motif DRACH (D = G/A/U, R = G/A, H = U/A/C). (F) Gene biotypes of DMP-associated genes. (G) Venn diagram showing the intersection of DMP-associated genes in each comparison. (H-J) Statistics of the number of DMPs on DMP-associated genes in trisomy 2L females compared with wildtype females (H), trisomy 2L males compared with wildtype males (I), and metafemales compared with wildtype females (J). CF, wildtype female control; CM, wildtype male control; 2LF, trisomy 2L female; 2LM, trisomy 2L male; XXX, metafemale; 5[1UTR, 5[1 untranslated region; 3[1UTR, 3[1 untranslated region.

**Figure 4—figure supplement 1.**
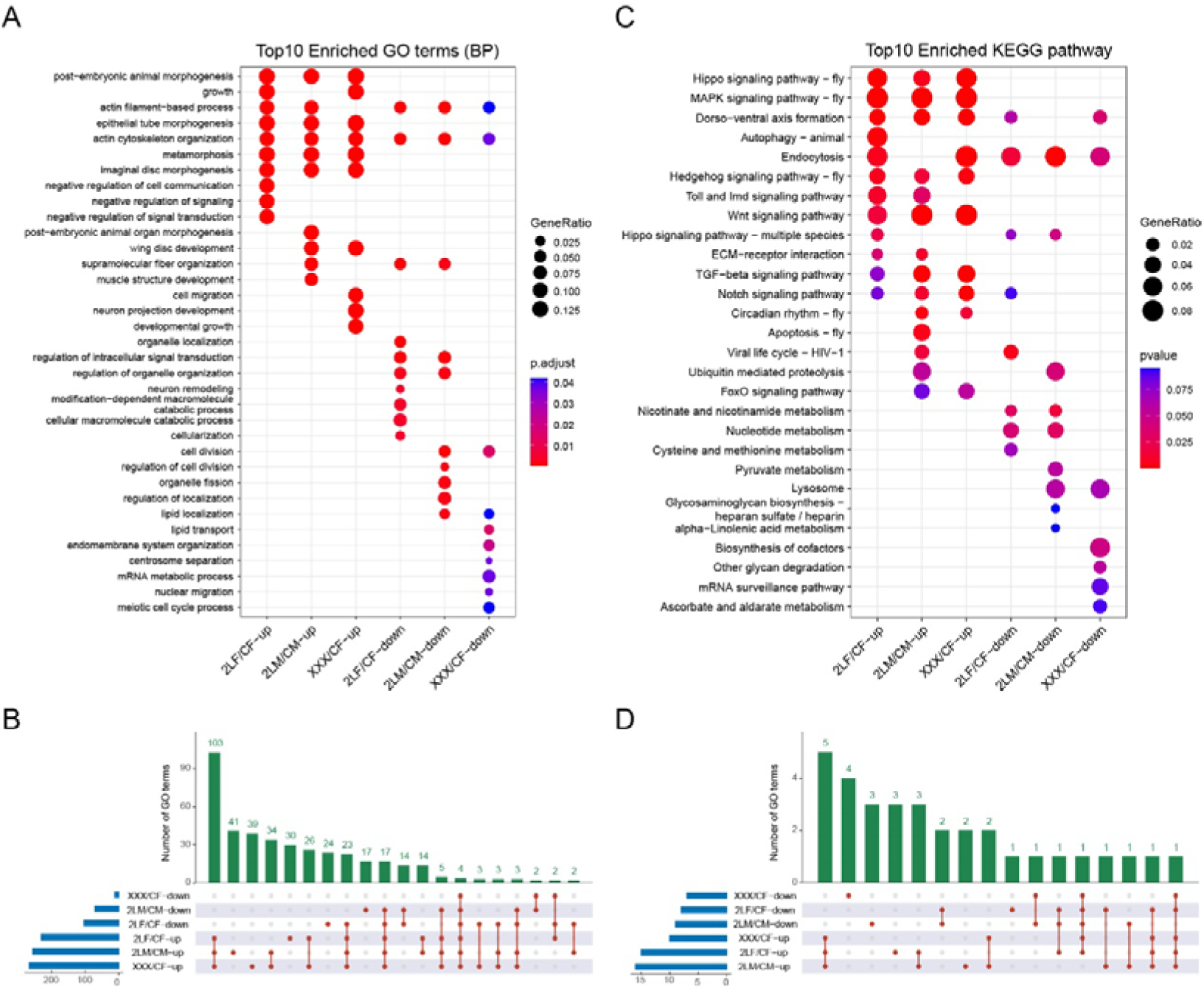
Functional and pathway analysis of DMP-associated genes. (A) Functional enrichment analysis of DMP-associated genes. Top 10 enriched GO terms (Biological Process) with adjusted p-value < 0.05 in each comparison were shown. (B) UpSet plot showing the shared sets of enriched GO terms of DMP-associated genes in different aneuploidies. (C) KEGG pathway enrichment analysis of DMP-associated genes. Top 10 enriched pathways with p-value < 0.1 in each comparison were shown. (D) UpSet plot showing the shared sets of enriched KEGG pathways of DMP-associated genes in different aneuploidies. CF, wildtype female control; CM, wildtype male control; 2LF, trisomy 2L female; 2LM, trisomy 2L male; XXX, metafemale.

**Figure 5—figure supplement 1.**
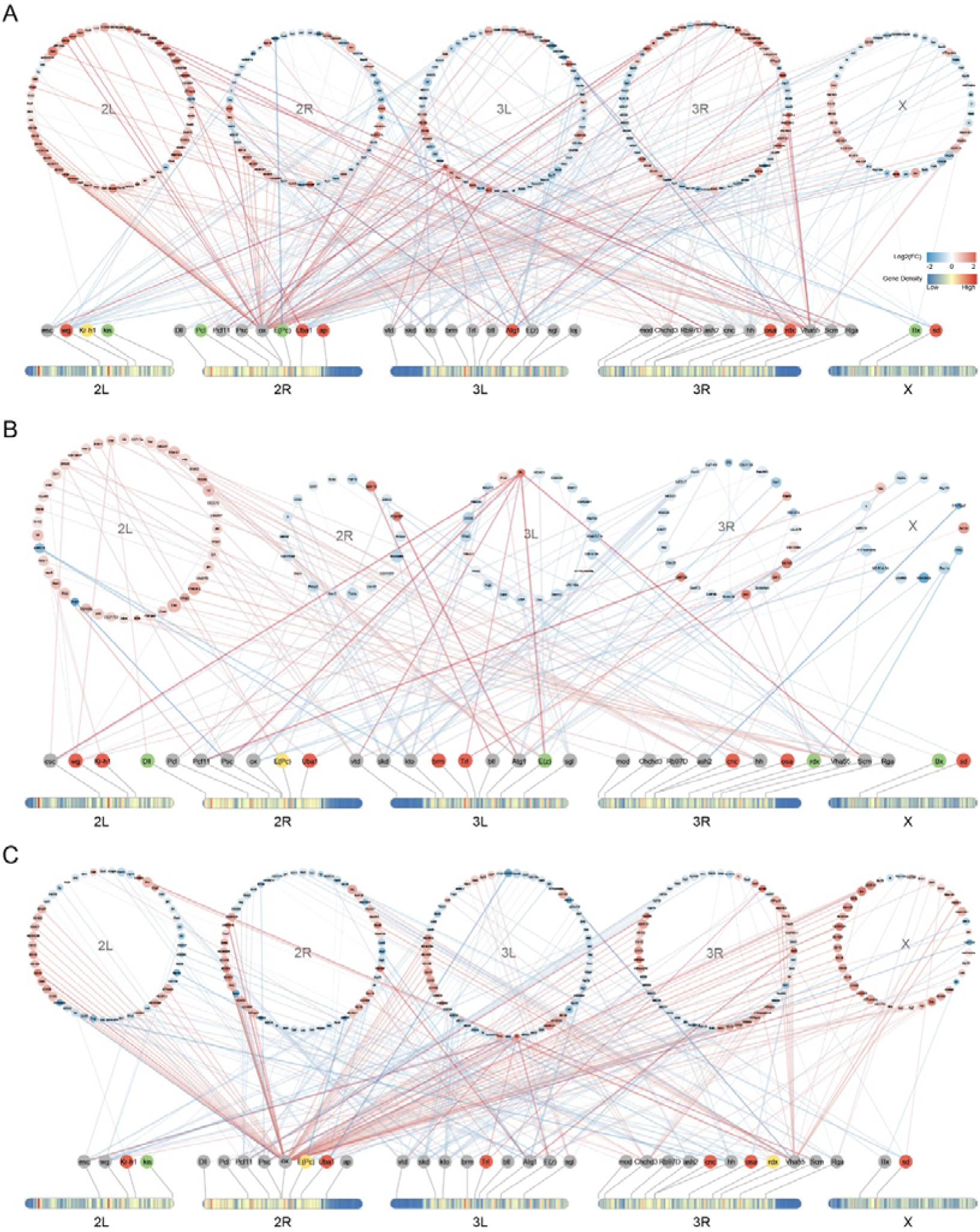
Dosage-sensitive modifiers and their interactors in aneuploid Drosophila. (A-C) Networks showing the methylation status of known dosage-sensitive modifiers and the expression changes of genes interact with these modifiers in trisomy 2L females (A), trisomy 2L males (B), and metafemales (C), respectively. Protein-protein interaction (PPI) relationships were obtained from the STRING database. The colors of the heatmaps on chromosomes represent the gene density. Dosage-sensitive modifiers were arranged according to their localizations, with red indicating genes with upregulated DMPs, green indicating genes with downregulated DMPs, yellow indicating genes with both up- and down-regulated DMPs, and grey indicating genes without DMPs. The interactors of these modifiers were also separated by chromosomes, and the colors of the nodes and edges indicate the log2(fold changes) of gene expression in trisomies. The width of the edges represents the combined score, and only interactions with combined score ≥ 700 were selected for presentation.

**Figure 6—figure supplement 1.**
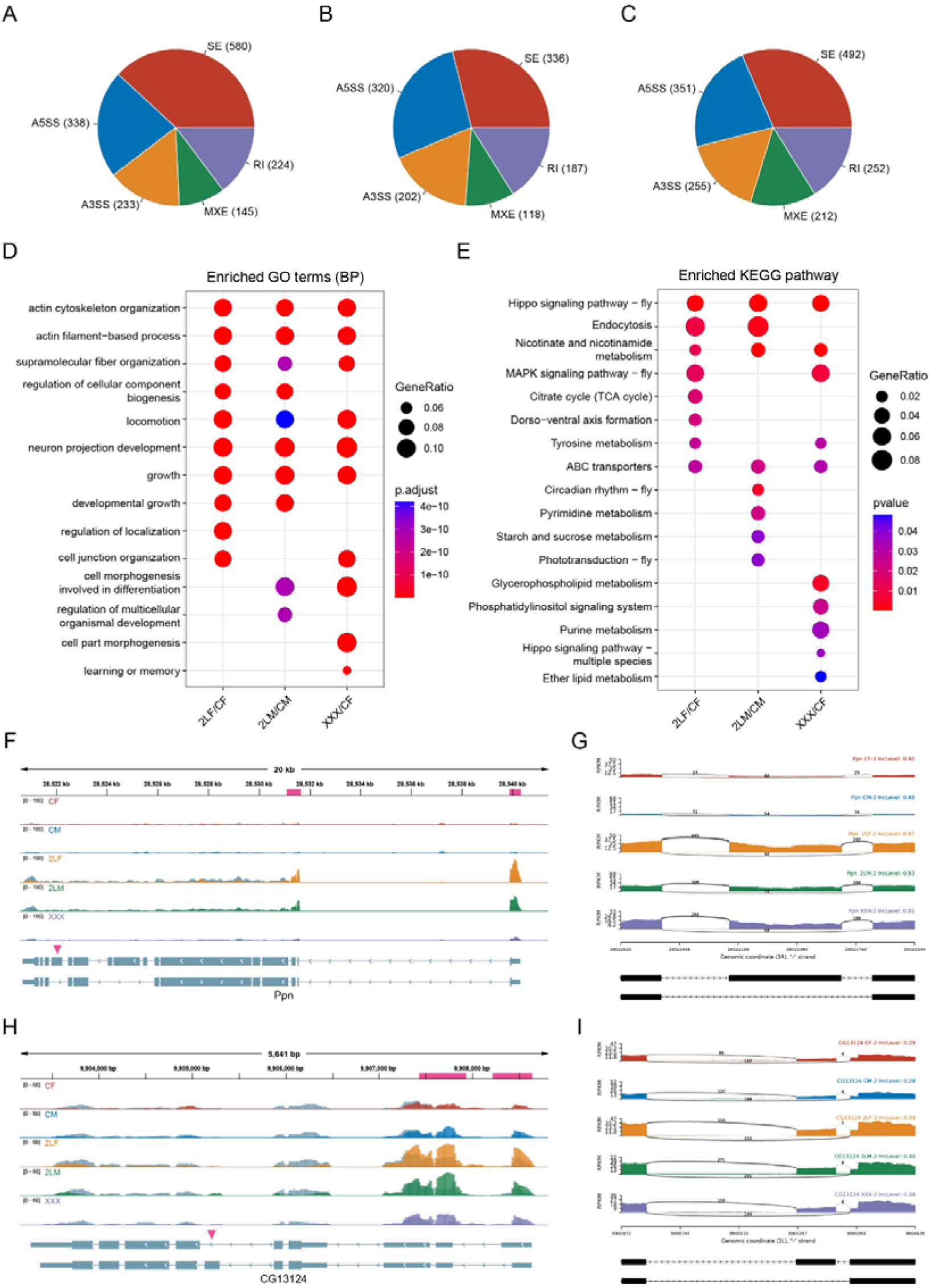
Alternative splicing analysis in aneuploidies. (A-C) Numbers of differential alternative splicing events in trisomy 2L females (A), trisomy 2L males (B), and metafemales (C). SE, skipped exon; A5SS, alternative 5’ splice site; A3SS, alternative 3’ splice site; MXE, mutually exclusive exons; RI, retained intron. (D) Functional enrichment analysis of differentially alternatively spliced genes. Top 10 enriched GO terms (Biological Process) with p-value < 0.05 in each comparison were shown. (E) KEGG pathway enrichment analysis of differentially alternatively spliced genes. Top 10 enriched pathways with p-value < 0.05 in each comparison were shown. (F,H) Genome browser example of Ppn (F) and CG13124 (H) for indicated MeRIP-seq data. Steelblue color represents input reads, while other colors represent IP reads. Signals were displayed as the mean CPM of two biological replicates. The gene architectures were shown at the bottom (only two representative transcript isoforms were shown). The magenta rectangles at above represent DMPs. The magenta arrowheads indicate the positions of differential alternative splicing. (G,I) Sashimi plots depicting RNA sequencing reads and exon junction reads at the positions where the differential splicing events occurs on Ppn (G) and CG13124 (I). The gene models were shown below. One of the biological replicates was chosen for representation. CF, wildtype female control; CM, wildtype male control; 2LF, trisomy 2L female; 2LM, trisomy 2L male; XXX, metafemale; CPM, Counts per million; RPKM, Reads per kilobase per million mapped reads.

**Figure 7—figure supplement 1.**
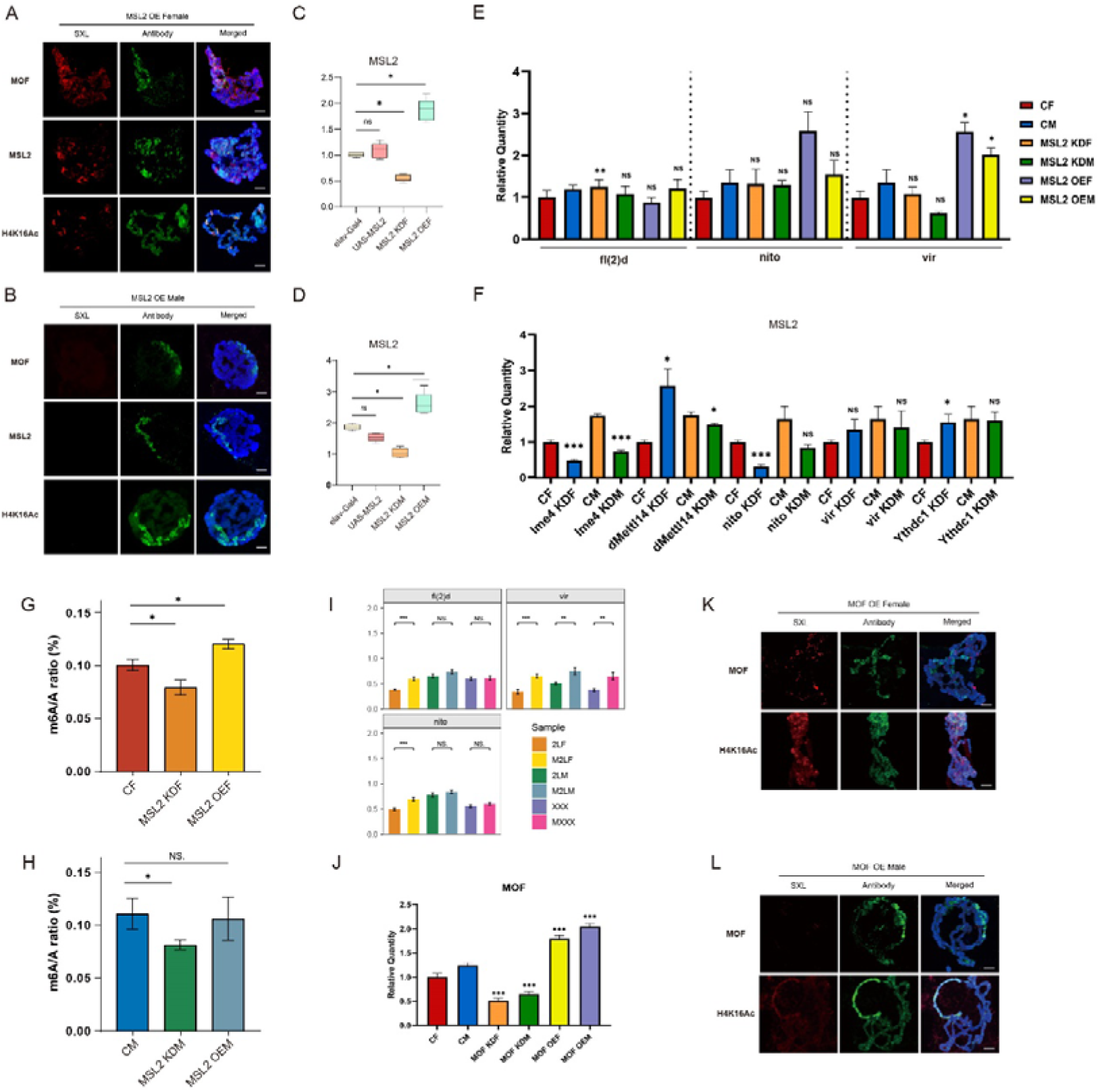
The relationships between m^6^A methylation and MSL complex. (A,C) Immunofluorescence of polytene chromosomes in MSL2-overexpressed female (A) and male (C) larvae. The red channel is the signal from SXL and the green channel is the signal from antibodies of MSL complex components. DNA is stained with DAPI in blue. Scale bars, 10 μm. OE, overexpressed. (B,D) RT-qPCR analysis of mRNA levels of MSL2 in the heads of MSL2 transgenic female (B) and male (D) Drosophila adults. Student’s t test *p < 0.05, **p < 0.01, ***p < 0.001. OE, overexpressed. (E) RT-qPCR analysis of mRNA levels of m^6^A regulators in the heads of MSL2 transgenic Drosophila. Student’s t test *p < 0.05, **p < 0.01, ***p < 0.001. CF, wildtype female control; CM, wildtype male control; OEF, overexpressed female; OEM, overexpressed male; KDF, knockdown female; KDM, knockdown male. (F) RT-qPCR analysis of mRNA levels of MSL2 in the heads of m^6^A regulators knockdown Drosophila. Student’s t test *p < 0.05, **p < 0.01, ***p < 0.001. CF, wildtype female control; CM, wildtype male control; OEF, overexpressed female; OEM, overexpressed male; KDF, knockdown female; KDM, knockdown male. (G,H) Abundance of total RNA m^6^A modification in the heads of MSL2 transgenic females (G) and males (H). Student’s t test *p < 0.05, **p < 0.01, ***p < 0.001. MSL2 KDF, MSL2 neural-knockdown female; MSL2 KDM, MSL2 neural-knockdown male; MSL2 OEF, MSL2-overexpressed female; MSL2 OEM, MSL2-overexpressed male. (I) RT-qPCR analysis of mRNA levels of m^6^A regulators in the brains of trisomy and MSL2-overexpressed trisomy Drosophila larvae. Student’s t test *p < 0.05, **p < 0.01, ***p < 0.001. 2LF, trisomy 2L female; 2LM, trisomy 2L male; XXX, metafemale; M2LF, MSL2-overexpressed trisomy 2L female; M2LM, MSL2-overexpressed trisomy 2L male; MXXX, MSL2-overexpressed metafemale. (J) RT-qPCR analysis of mRNA levels of MOF in the heads of MOF transgenic Drosophila. Student’s t test *p < 0.05, **p < 0.01, ***p < 0.001. CF, wildtype female control; CM, wildtype male control; OEF, overexpressed female; OEM, overexpressed male; KDF, knockdown female; KDM, knockdown male. (K,L) Immunofluorescence of polytene chromosomes in MOF-overexpressed female (H) and male (I) larvae. The red channel is the signal from SXL and the green channel is the signal from MOF or H4K16Ac. DNA is stained with DAPI in blue. Scale bars, 10 μm. OE, overexpressed.

**Figure 8—figure supplement 1.**
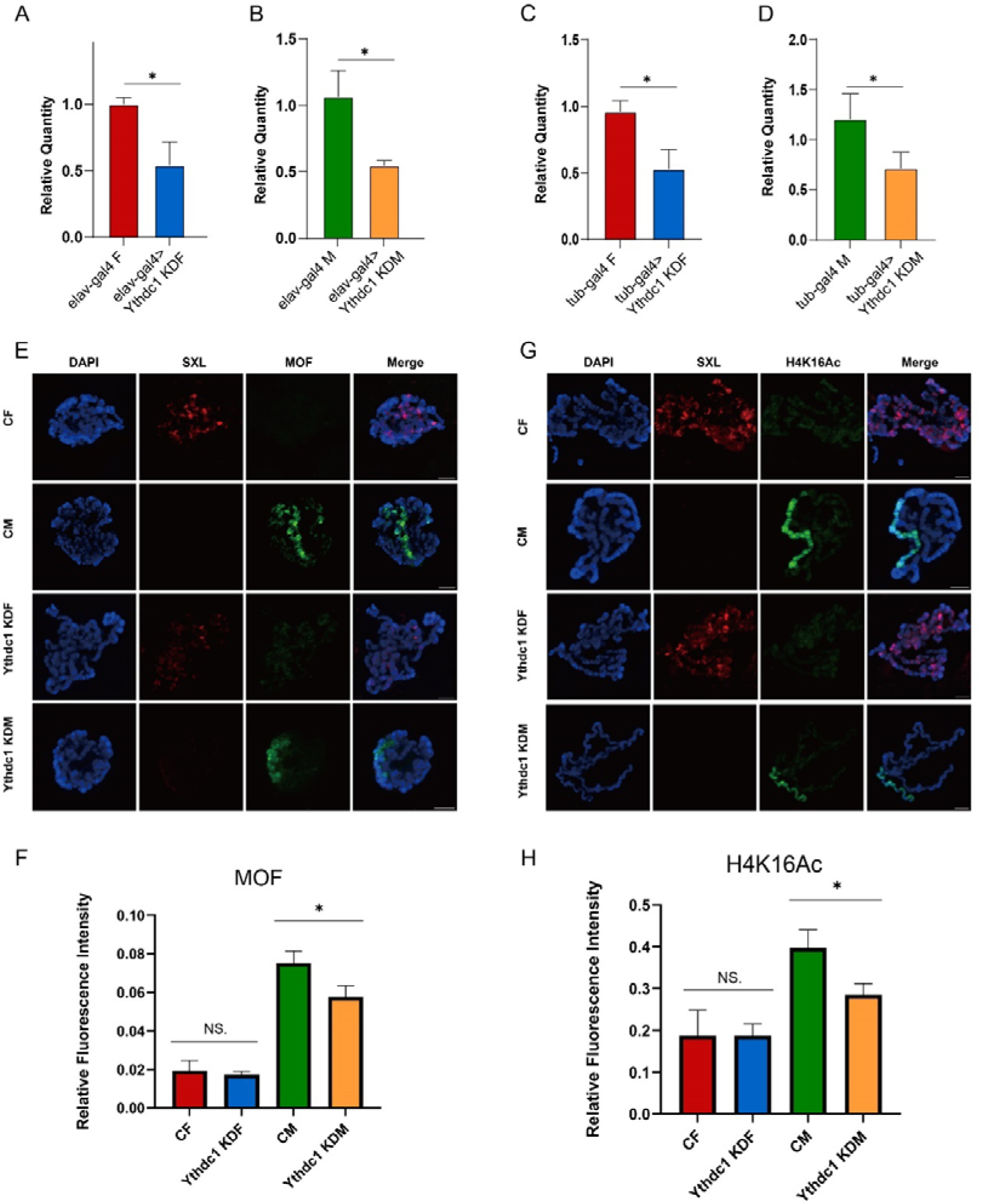
Ythdc1 may regulate H4K16Ac through histone acetyltransferase MOF. (A,B) RT-qPCR analysis of mRNA levels of Ythdc1 in the heads of Ythdc1 neural-knockdown female (A) and male (B). elav-gal4>Ythdc1 KDF, Ythdc1 neural-knockdown female; elav-gal4>Ythdc1 KDM, Ythdc1 neural-knockdown male. (C,D) RT-qPCR analysis of mRNA levels of Ythdc1 in Ythdc1 knockdown female (C) and male (D) adult Drosophila. tub-gal4>Ythdc1 KDF, Ythdc1 knockdown female; tub-gal4>Ythdc1 KDM, Ythdc1 knockdown male. (E) Immunofluorescence of polytene chromosomes showed the signals of SXL (red channel) and MOF (green channel) in the salivary glands of Drosophila larvae of different genotypes. (F) The levels of MOF in the salivary glands of Ythdc1 knockdown Drosophila quantified by immunofluorescence. (G) Immunofluorescence of polytene chromosomes showed the signals of SXL (red channel) and H4K16Ac (green channel) in the salivary glands of Drosophila larvae of different genotypes. (H) The levels of H4K16Ac in the salivary glands of Ythdc1 knockdown Drosophila quantified by immunofluorescence. Ythdc1 KDF, Ythdc1 knockdown female; Ythdc1 KDM, Ythdc1 knockdown male. Scale bars, 10 μm. Student’s t test *p < 0.05, **p < 0.01, ***p < 0.001.

**Figure 8—figure supplement 2.**
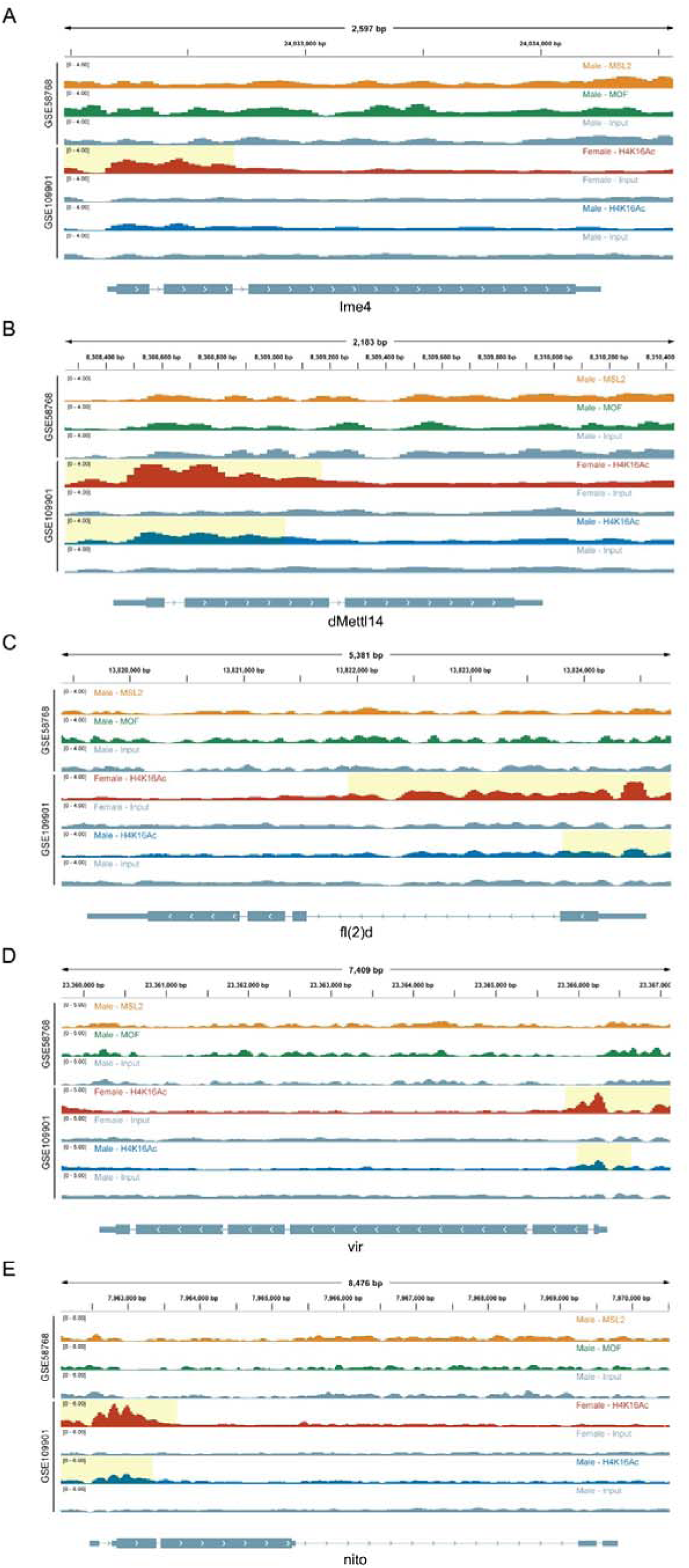
Genome browser example of m^6^A regulator genes for indicated ChIP-seq data. H4K16Ac, MSL2, and MOF ChIP-seq results for gene Ime4 (A), dMettl14 (B), fl(2)d (C), vir (D), and nito (E) are shown, respectively. Signals were displayed as CPM values. The gene architectures were shown at the bottom (only one representative transcript isoform was shown). The yellow shaded area indicates the presence of the H4K16Ac peaks.

